# The Three Waves: Rethinking the Structure of the first Upper Paleolithic in Western Eurasia

**DOI:** 10.1101/2022.10.28.514208

**Authors:** Ludovic Slimak

**Affiliations:** CNRS, UMR 5608, TRACES, Université de Toulouse Jean Jaurès, Maison de la Recherche, 5 Allées Antonio Machado, 31058 Toulouse, Cedex 9, France

**Keywords:** Grotte Mandrin, Ksar Akil, Neronian, Upper Paleolithic, Modern Human Expansion

## Abstract

The Neronian is a lithic tradition recognized in the Middle Rhône Valley of Mediterranean France now directly linked to *Homo sapiens* and securely dated to 54,000 years ago (ka), pushing back the arrival of modern humans in Europe by 10 ka. This incursion of modern humans into Neandertal territory and the relationships evoked between the Neronian and the Levantine Initial Upper Paleolithic (IUP) question the validity of concepts that define the first *H. sapiens* migrations and the very nature of the first Upper Paleolithic in western Eurasia. Direct comparative analyses between lithic technology from Grotte Mandrin and East Mediterranean archeological sequences, especially Ksar Akil, suggest that the three key phases of the earliest Levantine Upper Paleolithic have very precise technical and chronological counterparts in Western Europe, recognized from the Rhône Valley to Franco-Cantabria. These trans-Mediterranean technical connections suggest three distinct waves of *H. sapiens* expansion into Europe between 55-42 ka. These elements support an original thesis on the origin, structure, and evolution of the first moments of the Upper Paleolithic in Europe tracing parallel archaeological changes in the East Mediterranean region and Europe.

## Introduction

The recent attribution of the Neronian industry to *Homo sapiens* at around the 54^th^ millennium (56.8-51.7 ka cal. BP 95.4% prob.) at Grotte Mandrin in France not only indicates a 10,000-year push back of the arrival modern humans in Europe [1]; for the first time, concrete evidence of interactions between Neanderthals and modern populations are demonstrated in a specific territory. Five stratigraphic levels overlie the Neronian that have revealed Mousterian artifacts and Neanderthal teeth documenting the only occurrence of interstratification between modern and archaic hominins currently recognized in the world and a strict contemporaneity between these two populations. At Mandrin one year at most separates the preceding Neanderthal settlements and the arrival of modern humans (2-4], allowing us to approach the nature of potential interactions between these two populations. But the question of the first Upper Paleolithic (UP) is obviously not limited to only these exceptional archaeological records. This remarkable chronological and geographical overturning of our previously held theories about the first UP invites us to rethink the very structure of these human societies in Europe and, more broadly, in western Eurasia between 55 - 40 ka. This modern human incursion into Neanderthal territory and the relationships evoked between the Neronian and the Levantine Initial Upper Paleolithic – IUP-[1, 5, 6] question the validity of concepts that define the first *H. sapiens* migrations and the very nature of the first UP in western Eurasia. It is at this scale that the data from Mandrin invites rethinking the structure of the first UP, a period for which the most salient traits have stayed unchanged since the first half of the 20^th^ century.

This study discusses the structure of the connections that are now possible to establish between the banks of the eastern and western Mediterranean and underlines the unexpected technological connections and the remarkable cultural homogeneity of *H. sapiens* societies when they colonized Europe, which now seems likely to indicate the existence of three distinct waves of migration into the continent. I therefore hypothesize here that all of the first Upper Paleolithic industries recognized in the Ksar Akil sequence of coastal Lebanon have precise technical and chronological counterparts in Western Europe. I also posit that the almost unanimously hailed correlation that the Northern Early Ahmarian industries of Ksar Akil were a counterpart of the Protoaurignacian is false. The precise analysis of the technical successions of Ksar Akil allows us to defend a much broader position affecting the entire technical and historical structure of the first Upper Paleolithic and the articulations of a significant part of the so-called transitional industries of Western Europe.

The attribution of the Neronian to *H. sapiens*, its early chronology, and its insertion in the middle of the Western European Mousterian sequence raises fundamental questions about the anthropological framework that allows us to grasp such historical complexity. Among these questions immediately emerges inquiries about the origin of the Neronian. Apart from Grotte Mandrin, few other Neronian sites exist (Moula IV, Néron I, Maras, and Figuier 1 and 1’), all having small assemblages that were excavated long ago with pickaxes [7, 8], and all occurring within a restricted portion of the middle Rhône valley.

However, locally and diachronically, a rather remarkable technological continuity is documented in the Mandrin sequence between the Neronian and overlying Protoaurignacian, in Levels E and B1 respectively. Technologically, this process of continuity is clearly marked and could be summarized by the change from the use of hard (mineral) to softer organic percussion in the production of lithic artifacts [5-10] and from the use of a facetted striking platform to one that is straight and abraded. Thus, no other technological peculiarity makes it possible to fundamentally distinguish these two sets; they are remarkably similar in their technological structures, their production objectives, specific features of the transformation of the materials involving ventral and alternating retouch created by pressure retouching into the palm, and the function of the obtained tools. The Neronian, carried by *H. sapiens*, can therefore be understood technically and historically as a Pre-Protoaurignacian or a Protoaurignacian 0. There is however no evidence for technological continuity between the underlying Rhodanian Quina Mousterian from Level F and the Neronian from Level E, the only strata where the two industries are superimposed and stratigraphic mixing can be excluded (Supplementary Note 1). This proposition questions the origins of the Neronian and Protoaurignacian, and the processes documenting the structuring of these technical traditions. While we note connections between the Neronian and the Protoaurignacian, precise modalities of such evolution remain unclear and no local origin for Neronian can be discerned from the middle Rhône valley.

### Rethinking of the origin of the Upper Paleolithic: a Mediterranean Odyssey

The Ksar Akil sequence occupies a key position in the understanding of Paleolithic societies in the eastern Mediterranean. The site is located 10 km northeast of Beirut and overlooks the coastal plains on the foothills of Mount Lebanon. Numerous archaeological excavations have been undertaken there, revealing 22.6 m of archaeological deposit from the Middle Paleolithic to the Epipaleolithic, and the site constitutes one of the most complete records currently recognized in Eurasia regarding the transition between the MP and UP. These archaeological levels were reached during two excavation phases held in 1937-1938 and 1947-1948 led by Ewing [11, 12]. Tixier’s operations from 1969 to 1975 encountered only the upper part of the sequence and a few subsequent phases of the UP that are not directly relevant here [13]. The sequence has been subdivided into 36 main archaeological units; I restrict my discussion here to the 31 MP and UP archaeological units. From to base to summit:

- Levels XXXVI to XXVI are from the Middle and late MP;
- Levels XXV to XXI involve the IUP;
- Levels XX and XVI relate to the initial phases of the Early Upper Paleolithic.

These stratigraphic successions form part of a unique context where technological and biological aspects of the origin of the UP can be concretely addressed [14, 15].

Connections between the European and the Mediterranean Levantine archaeological records have been considered since the early 20^th^ century. When it was recognized that the Aurignacian represented the first European UP, the same Aurignacian was simultaneously in the Levant area [16-17], as exemplified when during a conference in London in 1969, Bordes employed the term “Aurignacian in its strict sense” for the assemblages from Ksar Akil’s levels IX and X of the 1937-38 excavations (Xc-Xia levels of the 1947-48 excavations) [13, 18]. If, in Europe, a component prior to this form of the Aurignacian was proposed in the 1960s [19-21], its many detractors questioned the very existence of such industries until the almost definitive abandonment of these ideas at the end of the 1970s [22]. This hypothesis did not gain momentum again until the turn of the 1990s [23-26]. The recognition of this Protoaurignacian and its chronological position prior to the early Aurignacian finally helped produce correlations between the eastern and western shores of the Mediterranean. These quickly became formalized, suggesting the existence of a strict technological and cultural unity between the European Protoaurignacian and the Levantine Early Ahmarian [27-32]. These correlations were again based largely on the Ksar Akil reference sequence, the only to document all phases of the first UP in the Eastern Mediterranean (Supplementary Note 2), but one excluded from the most recent series of formalized comparisons between Levantine and European UP assemblages [33-35]. These historiographic details are essential to understand the viability of these proposed connections across the Mediterranean.

The use of the term IUP groups together collections from varied origins recognized over a vast territory ranging from North Africa to the highlands of Central Asia, and therefore encompasses quite diverse technical realities. In this regard, it is necessary to differentiate the generic term IUP, which does not have precise techno-cultural value [36], from the IUP of Ksar Akil which refers to a very precise technical reality. And because of Ksar Akil’s place in the history of research, it should be used as a type-sequence for the determination of an IUP *stricto sensu*, as compared to a IUP *lato sensu*, a name therefore grouping together a large fraction of the first UP industries of the Old World with no suggestion of a precise technical or cultural connection. My use of the term IUP hereafter is *stricto sensu*, as it is documented in the Ksar Akil sequence.

To position Mandrin’s archaeological record in the larger Eurasian context, analyses of Ewing’s 1947-1948 collections at Harvard University’s Peabody Museum of Archaeology and Ethnology were undertaken from 2016 to 2019. The stratigraphic subdivisions of this specific collection are notably more precise than from the 1937-1938 excavations (which are mainly curated by the British Museum in London), and include the full stratigraphic sequence, therefore making this collection the most important here. For example, layer IX from the 1937-1938 excavation, nearly 2 meters thick, was subdivided into 6 subunits (a-f) during the 1947-1948, detailing technological changes recorded between each archaeological unit. Although not corresponding to current standards, great attention was nevertheless paid regarding the smallest archaeological elements during excavation [37]. Compared to the 1937-1938 series which was relocated various times resulting in loss of part of the collection, Harvard’s 1947-1948 collection has been much less handled [37]. Despite being the most precise stratigraphically and the most accurate regarding excavation methods, the Harvard collection has been less studied.

This research focused on 31 units, ranging from the MP to the first UP, levels XXXVI to XIII. A total of 17,809 lithic pieces were analyzed and integrated into a database distinguishing 138 distinct technical and typological categories to account for the main specificities of these industries. These collections were also photographed, technically drawn, and functionally analyzed by Laure Metz (U. Connecticut, UMR LAMPEA). The aim of presenting the elements here is to put these data into qualitative perspective with what has been proposed regarding relations between Europe/Levant, essentially on bibliographic bases, concerning the first moments of the UP. This presentation therefore focuses primarily on layers XXV to XIII. The qualitative analysis gives a clear impression of continuity within these 13 stratigraphic units, illustrating technical evolution expressed gradually from one unit to another, as noted by Ohnuma and Bergman [38]. Based on technical systems, it thus seems impossible to distinguish a IUP unit superimposed by a completely EUP unit. However, technical peculiarities appear very clearly if we compare the XXIII-XXII units of the IUP with the XVI-XVII units of the EUP. The low artifact count (n=33) and combination of typical MP and UP technologies in layer XXV suggest that it is the product of mixing during excavation.

The image that emerges from these quantitative analyses of Ksar Akil’s technical record is that of an abrupt break between layers XXVI and XXIV, precisely between the MP and UP assemblages (Figs. 1-6). We can deduce that not only no serious mixing between stratigraphic units can be documented but also that, from the point of view of technical lithic systems, the sequence locally shows no possibilities of continuity between the MP and UP. At the same time, the technical breaks visible between the end of the MP and the beginning of the UP mean that the question of the emergence of the IUP cannot be documented from this sequence. These data suggest either that Ksar Akil presents an absence of archaeological deposit over a relatively long period of time, separating units XXVI and XXIV (the time required for the emergence of the IUP from a local technical substrate), or that the IUP was intrusive in this region [39-40]. If the Ksar Akil sequence can be considered fundamental to understanding the beginnings of the UP in Eurasia, the sequence, however, may not record all of the phases of its development. In this geographic area, sequences like Boker Tachtit may illustrate some of the first phases of this emergence. These early stages could have been structured around obtaining massive points from bipolar debitage, the technical affinities of which with the Bohunician of Central Europe have already been noted [33-35]. These data suggest the possibility of a fourth technical time, prior to the oldest phases recorded at Ksar Akil, putting into question the origin, in time and space, of the points systems at the beginning of the UP, whose source could be sought after more broadly in the geographic areas between the Mediterranean Levant and Central Asia [36, 41]. At the same time, within the Ksar Akil type sequence, we immediately notice that the clear processes of continuity that we evoked from the IUP to the EUP are inscribed here in concrete technical and stratigraphic realities. This should make it possible to understand, with unique resolution on the scale of Eurasia, the structure of the first moments of the UP and the evolution of these technical processes over time.

**Fig. 1.**
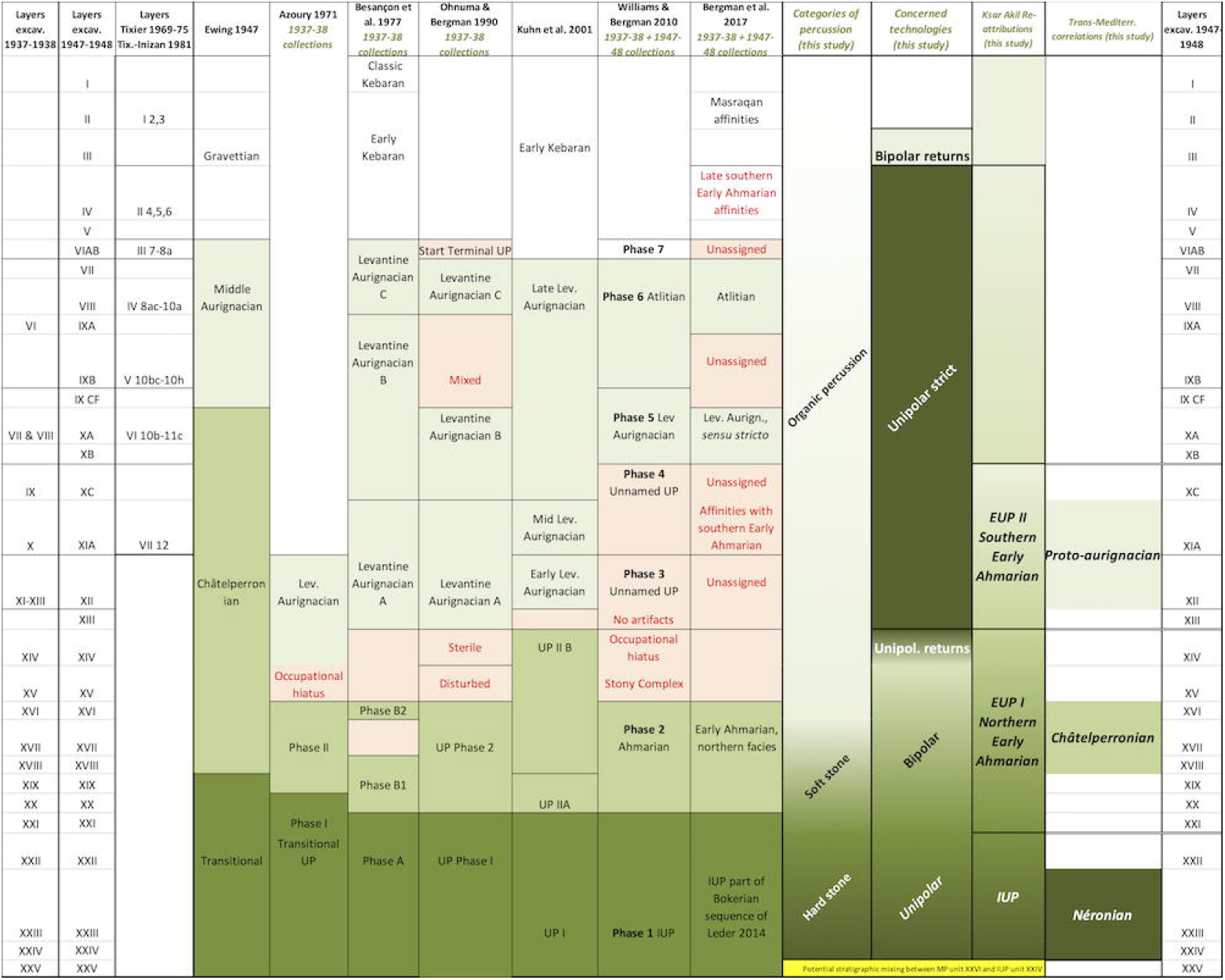
Summary of interpretations from the Ksar Akil sequence from 1947 to 2017 (11, 37, 38, 52, 77-79). The columns on the right present the keys to the technical and cultural readings based on my analyses and interpretations.

**Fig. 2.**
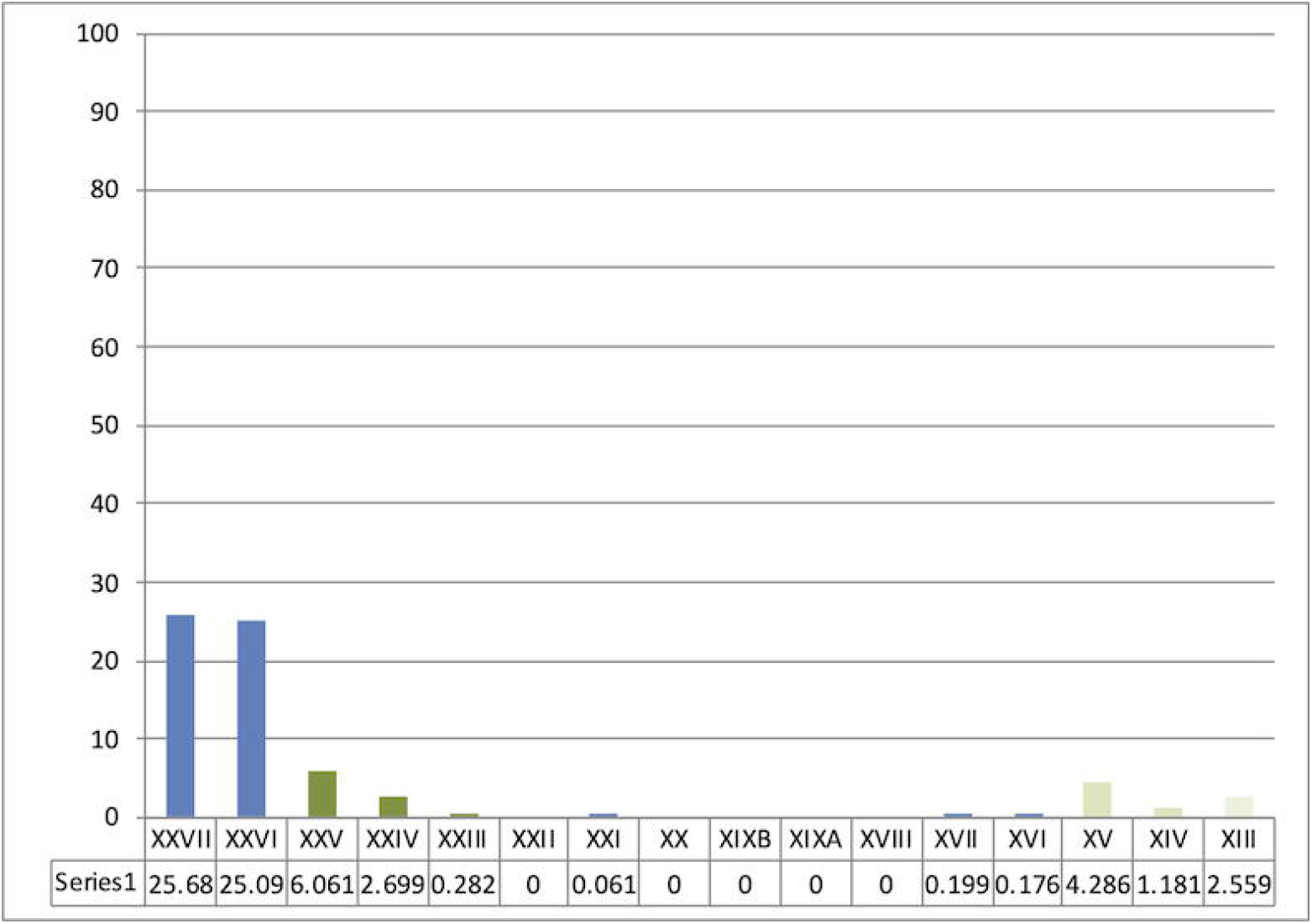
Sequence from Ksar Akil, 1947-1948 excavations. Representation of Levallois debitage between the Middle Paleolithic units, layers XXVII-XXVI, and the sequence of the beginnings of the Upper Paleolithic until layer XIII. Even though Levallois debitage represents more than 25% of the assemblage in the last units of the Mousterian (blue), they are virtually absent or anecdotal from the very start of the Upper Paleolithic (green). These representations illustrate a clear and abrupt rupture between the Middle and the Upper Paleolithic. Layer XXV, the first IUP unit, documents the highest proportion of these Mousterian debitages. This XXV unit is only composed of a few lithic pieces and this lithic assemblage could well be artificial and only constitute a mix of layers XXVI and XXIV.

**Fig. 3.**
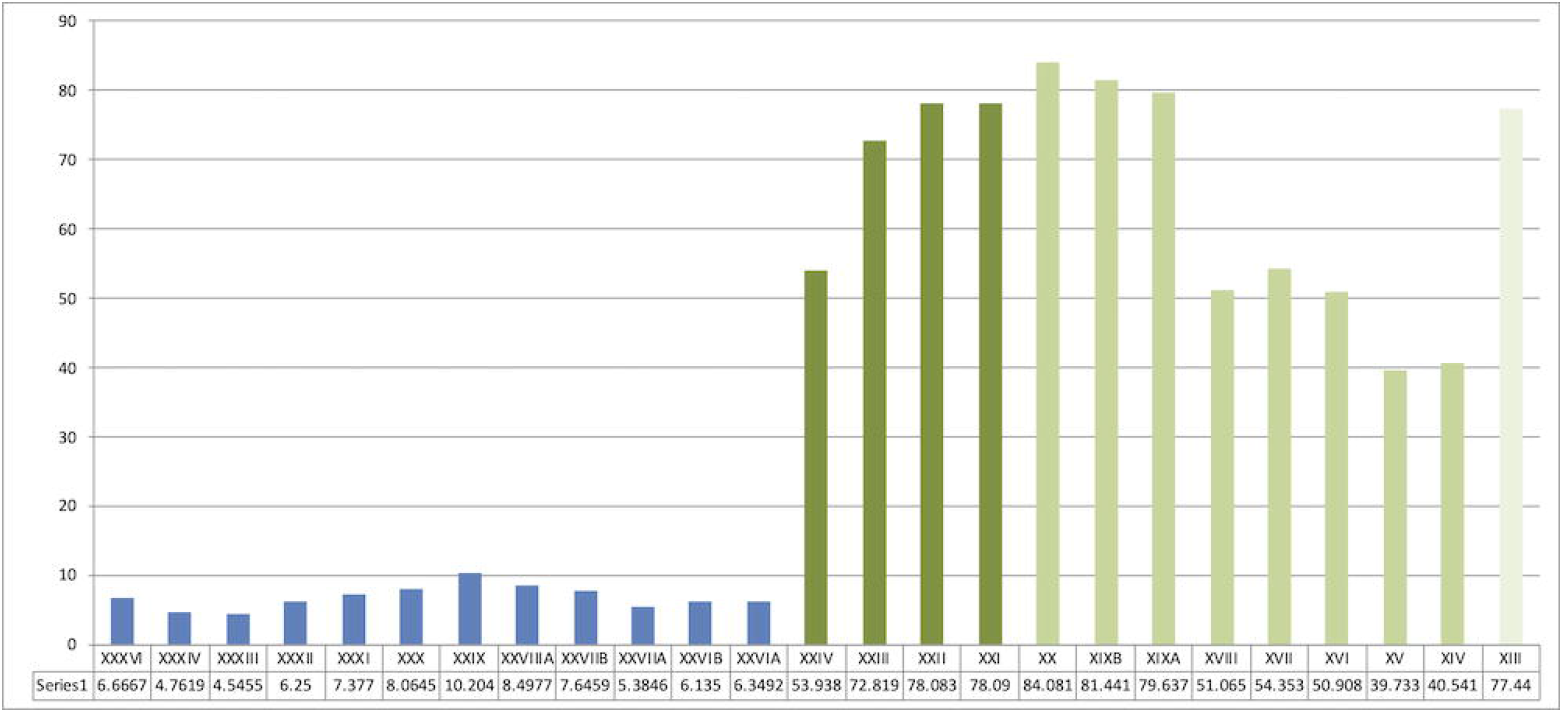
Sequence from Ksar Akil, 1947-1948 excavations. Representation of laminar blanks (blades and bladelets) and points within the Mousterian sequence (blue) and the first three phases of the Upper Paleolithic (green). Blades and points abruptly appear in the sequence with no possibility of continuity between the end of the Mousterian and the IUP.

**Fig. 4.**
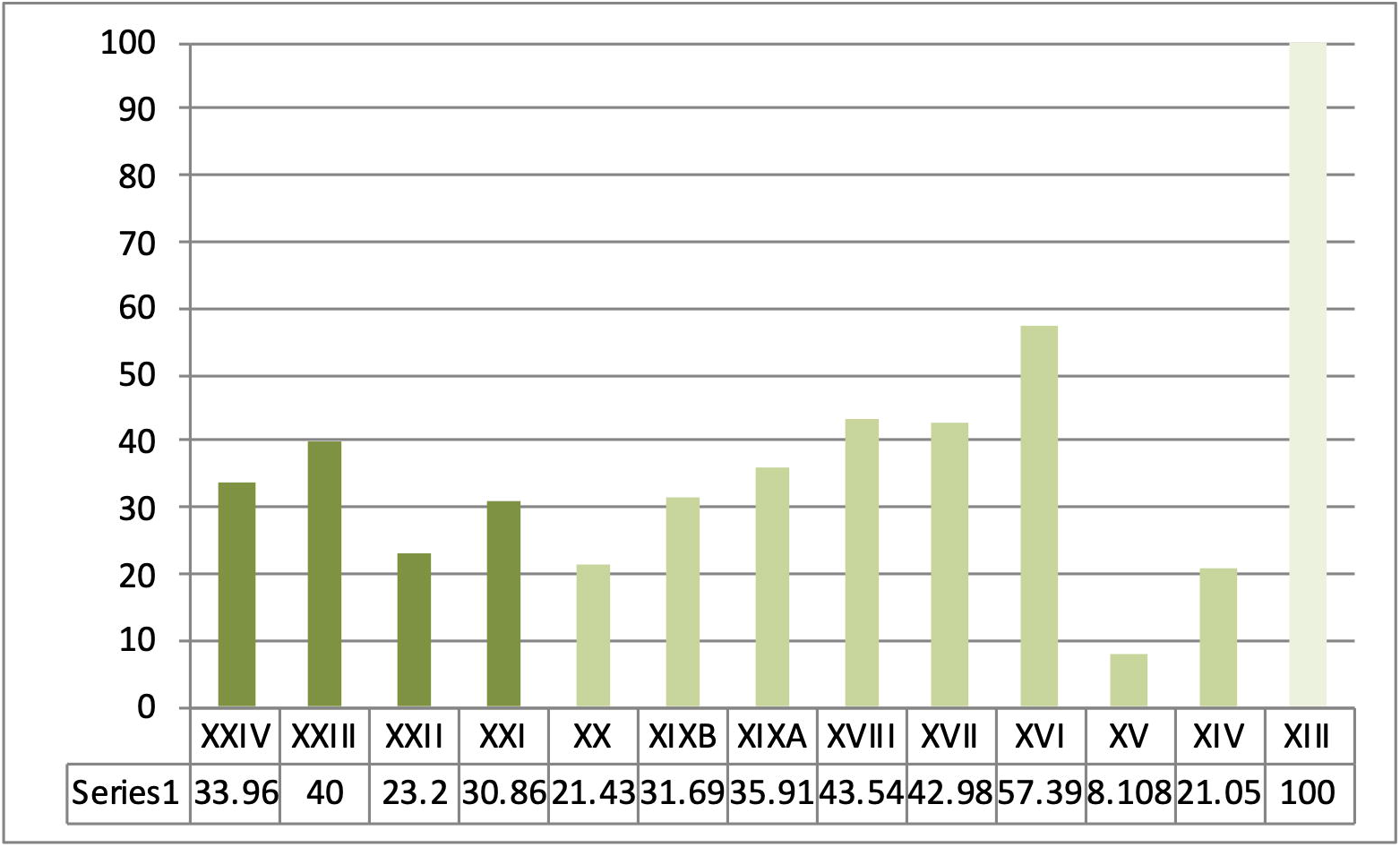
Sequence from Ksar Akil, 1947-1948 excavations, located at the Peabody Museum, Harvard. Representation of microlith products -bladelets and micropoints-in the first phases of the Upper Paleolithic, IUP (dark green), EUP I/NEA (medium green), and EUP II/SEA (light green).

**Fig. 5.**
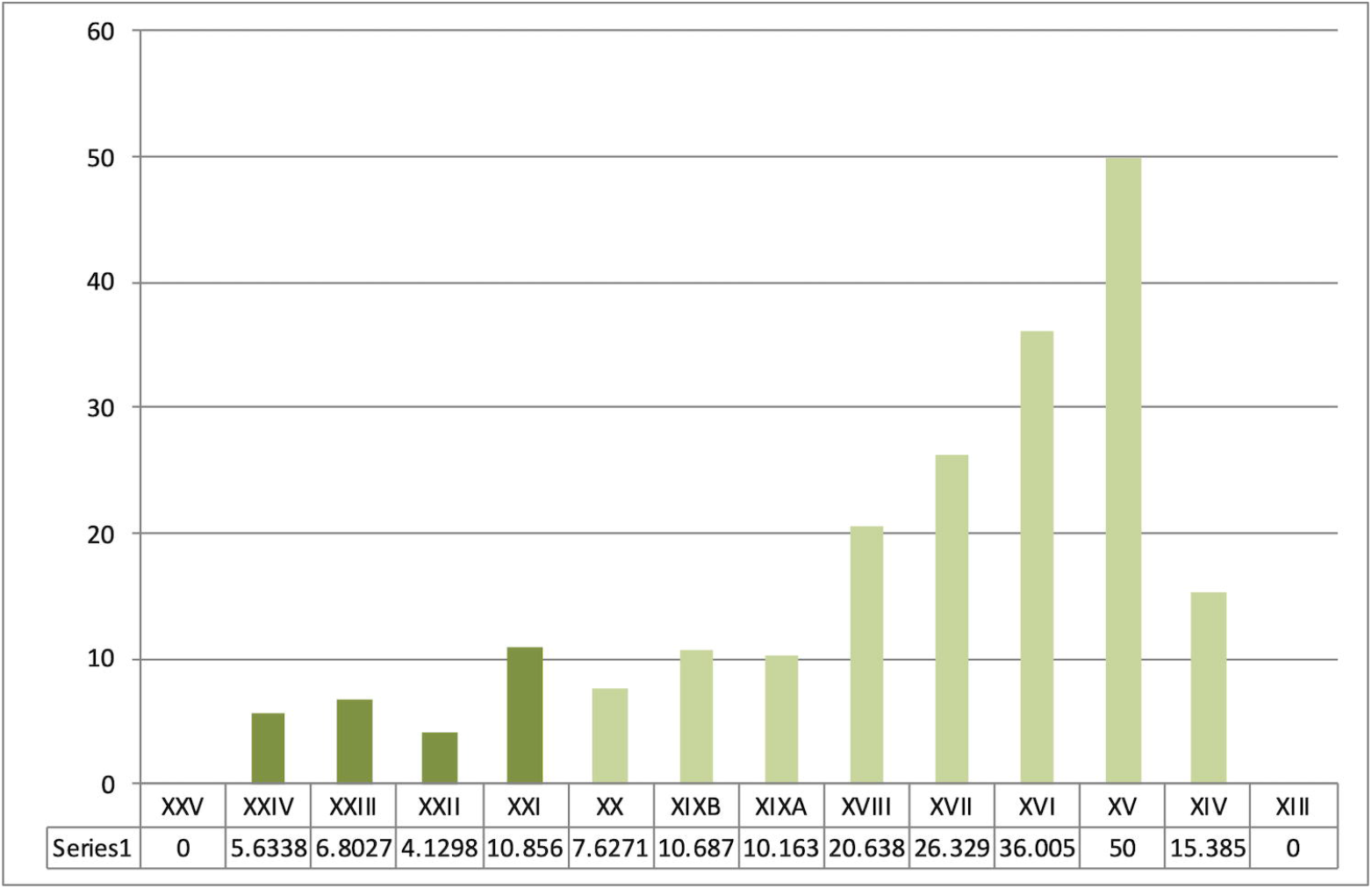
Sequence from Ksar Akil, 1947-1948 excavations. Representation of bipolar productions within the blade and point debitage of the IUP (dark green), the EUP I/NEA (medium green), and the EUP II/SEA (layer XIII, 0%).

**Fig. 6.**
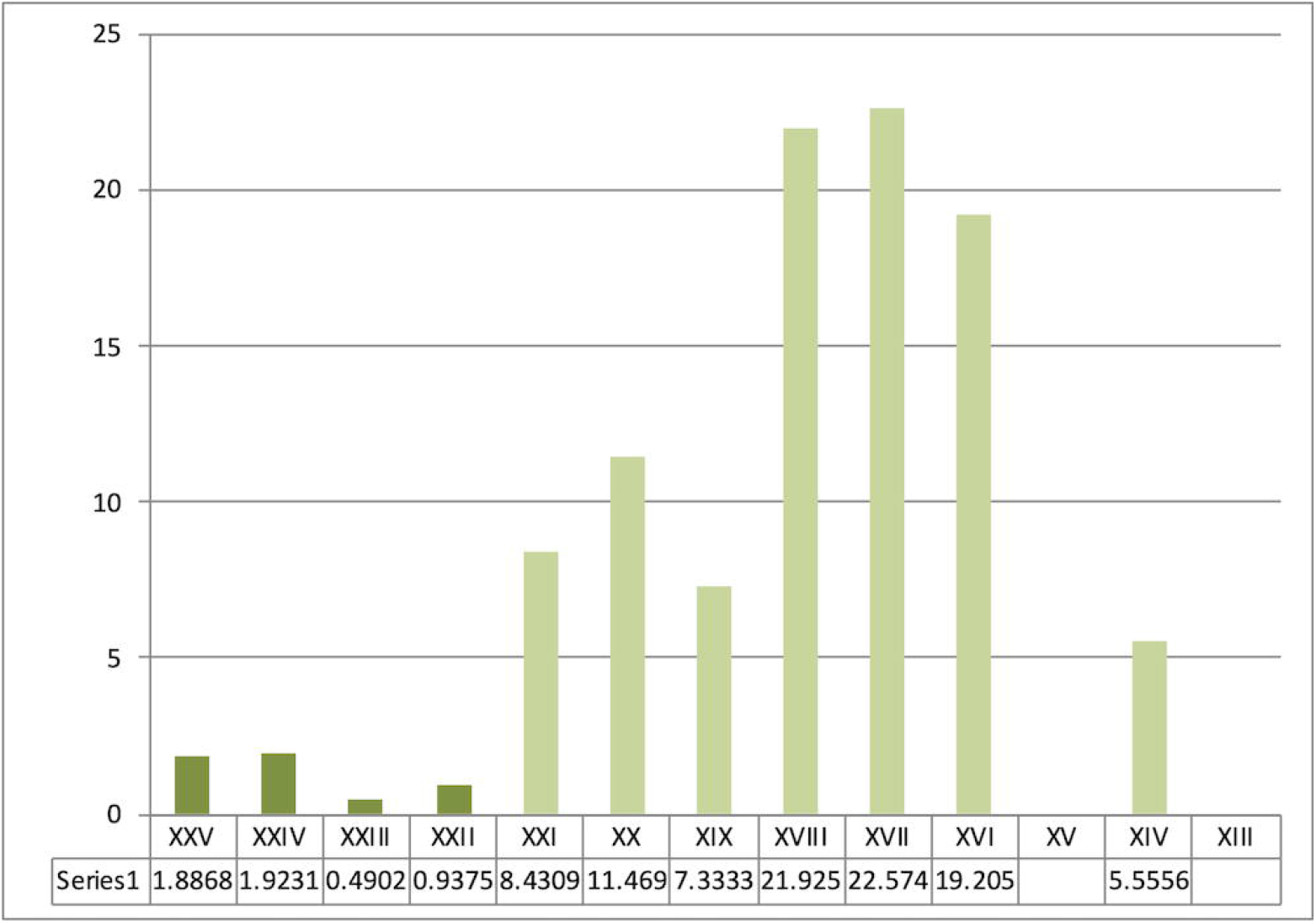
Representation of backed retouched tools from Ksar Akil within the typological corpus in the IUP (dark green), EUP I/NEA (medium green), and EUP II/SEA (layer XIII, 0%). (a) 1937-1938 excavations (British Museum, Ohnuma 1988). (b) 1947-1948 excavations.

### Back to Mandrin

Analysis of the Ksar Akil industries from Father Ewing’s excavations allows us to recognize the existence of three distinct phases at the turn of the UP. This phasing only partially overlaps with previously proposed frameworks, particularly concerning the last moments of the Northern Early Ahmarian (NEA) and its relationship with the overlying industries. In any case, three significant phases can be clearly distinguished, beyond the processes of continuity in the structure of the technical systems that can be highlighted from layers XXV to XIII at Ksar Akil (Supplementary Note 4);

- a first phase, of unipolar Levallois points (IUP; Fig. 7);

**Fig. 7.**
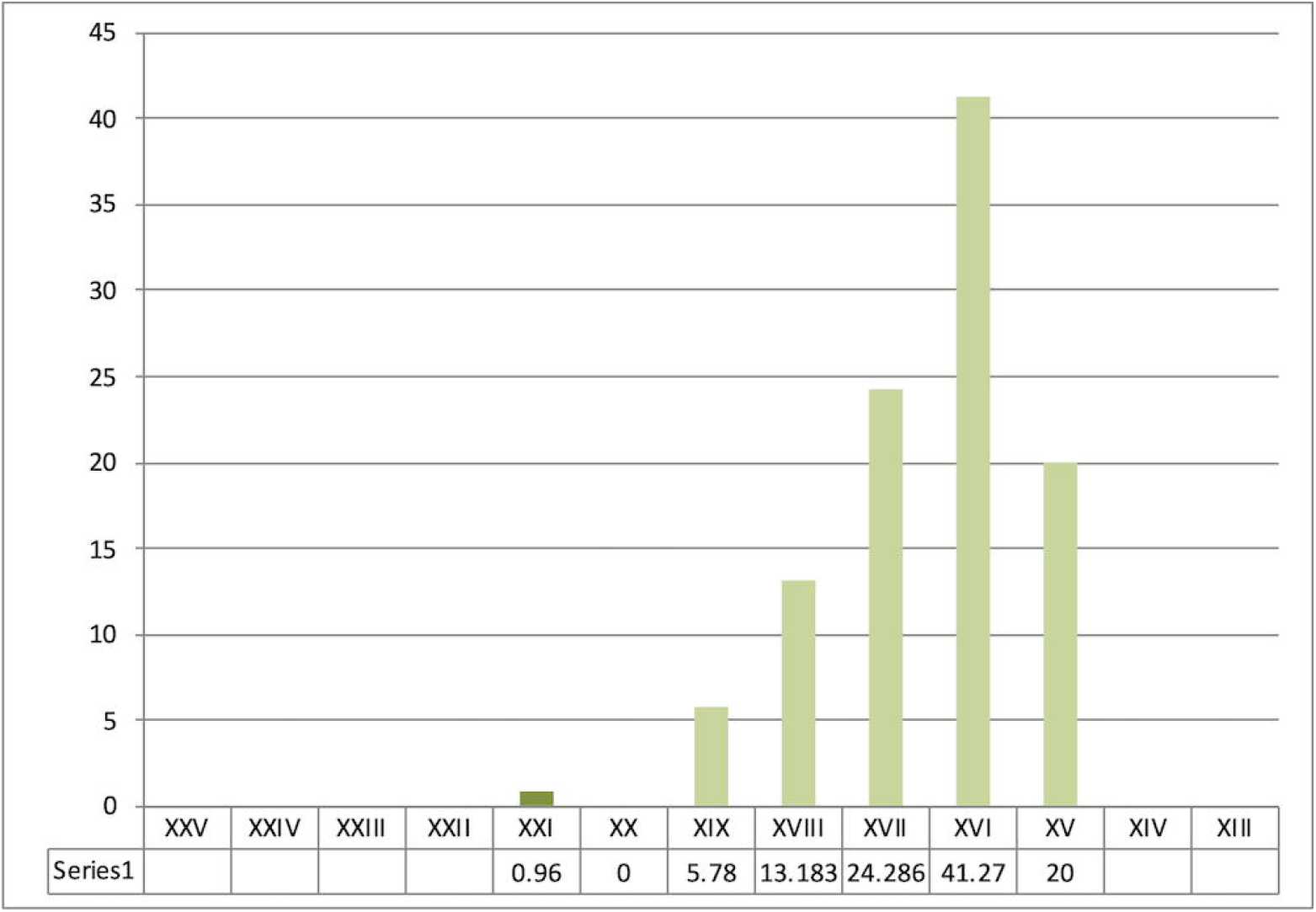
Sequence from Ksar Akil, 1947-1948 excavations, located at the Peabody Museum, Harvard. Points and blades from the Initial Upper Paleolithic of Ksar Akil, layers XXV-XXII. Drawings by L. Metz.
- a second phase, of backed points mainly from bipolar laminar debitages (EUP/ NEA);
- a third phase, of rectilinear acute bladelets issued from unipolar convergent debitages (layers XIII and above).

Unanimously used for close to 20 years, the correlation between this second phase of Ksar Akil’s Early Ahmarian and the Protoaurignacian [28, 32, 42-46] must be abandoned definitively. These systems do not overlap technologically, technically, nor typologically.

Clear East/West correlations can however be established between the Levant and Western Europe. In this correlation, an equivalent of the Protoaurignacian can easily be recognized in the layer XIII industries. This proposal here is close to the conclusions of Kadowaki *et al*. [47], but they proposed correlations with Ksar Akil layers IX-XI and then with more recent units from the Ksar Akil sequence. They also proposed that the Southern Early Ahmarian (SEA) was more recent than the NEA, the two having no stratigraphic overlay, and that the Protoaurignacian chronologically preceded the SEA. I propose here that, from the point of view of the general technical structure of these industries, these layers IX-XI do not represent the oldest industries of Ksar Akil that are technically comparable to the Protoaurignacian, which I place as early as layer XIII (Supplementary Notes 2 & 4).

Here I propose that prior to phase 3/SEA/Protoaurignacian of this chrono-cultural breakdown of Ksar Akil, the two other technical phases of this sequence also have direct parallels in the European records. The Neronian, entirely based on the production of unipolar convergent points and micropoints, technically represents a perfect replica of the Levantine phase 1/IUP. The technical systems, the production objectives, the morphology, and even the morphometry of the sought-after points are strictly identical [1, 5, 6]. In parallel, the function of the points, determined in functional analysis by Laure Metz, shows that the points of Ksar Akil XXV-XX and those of Mandrin E fall strictly within the same functional categories [48-49]. In both cases, they are mainly projectile points used with mechanical propulsions -spearthrower and/or bow. Morphometric width/thickness analysis shows that no differentiation can be made between Neronian and IUP points (Fig. 8). No distinction can be made here between these technical systems, even though they are located at opposite ends of the Mediterranean. We have also seen that, although radiometric approaches in the Levant still provide disputable results, there is every reason to believe that the beginnings of the Levantine IUP are contemporary with the Neronian of Mandrin. We also know that the Neronian was created by *H. sapiens* populations who were exotic to this region and who settled for some time in Neandertal territory [1]. All these data allow us to deduce that the two cultural groups, the Levantine IUP *stricto sensu* (as recognized in Ksar Akil) and the Neronian, actually are one. The question of the chronology of Ksar Akil’s IUP has produced clearly divergent models, but the data here would be compatible with the model proposed by Bosch [50, 51] who concluded that the ages obtained from the IUP represent minimum ages (Supplementary Note 3). Although the actual age of the beginnings of the IUP at Ksar Akil is still unknown, Bosch dated layer XXII to >46 ka and the IUP begins appearing in layer XXV, thus substantially older than 46 ka. If we widen the focus to other types of evidence, the presence of shells for example, numerous at Ksar Akil, but absent in Mandrin E, does not allow us to individualize the Neronian from the IUP, seeing that the shells of mollusks in the IUP, pierced or not, are almost exclusively recognized at coastal sites [52, 53]. The transformation of bones or teeth to produce objects of symbolic value is also well attested in the Neronian [1]. The evidence of a *H. sapiens* tooth in the Neronian also supports the correlations between the Neronian and the IUP presented here [5, 6].

**Fig. 8.**
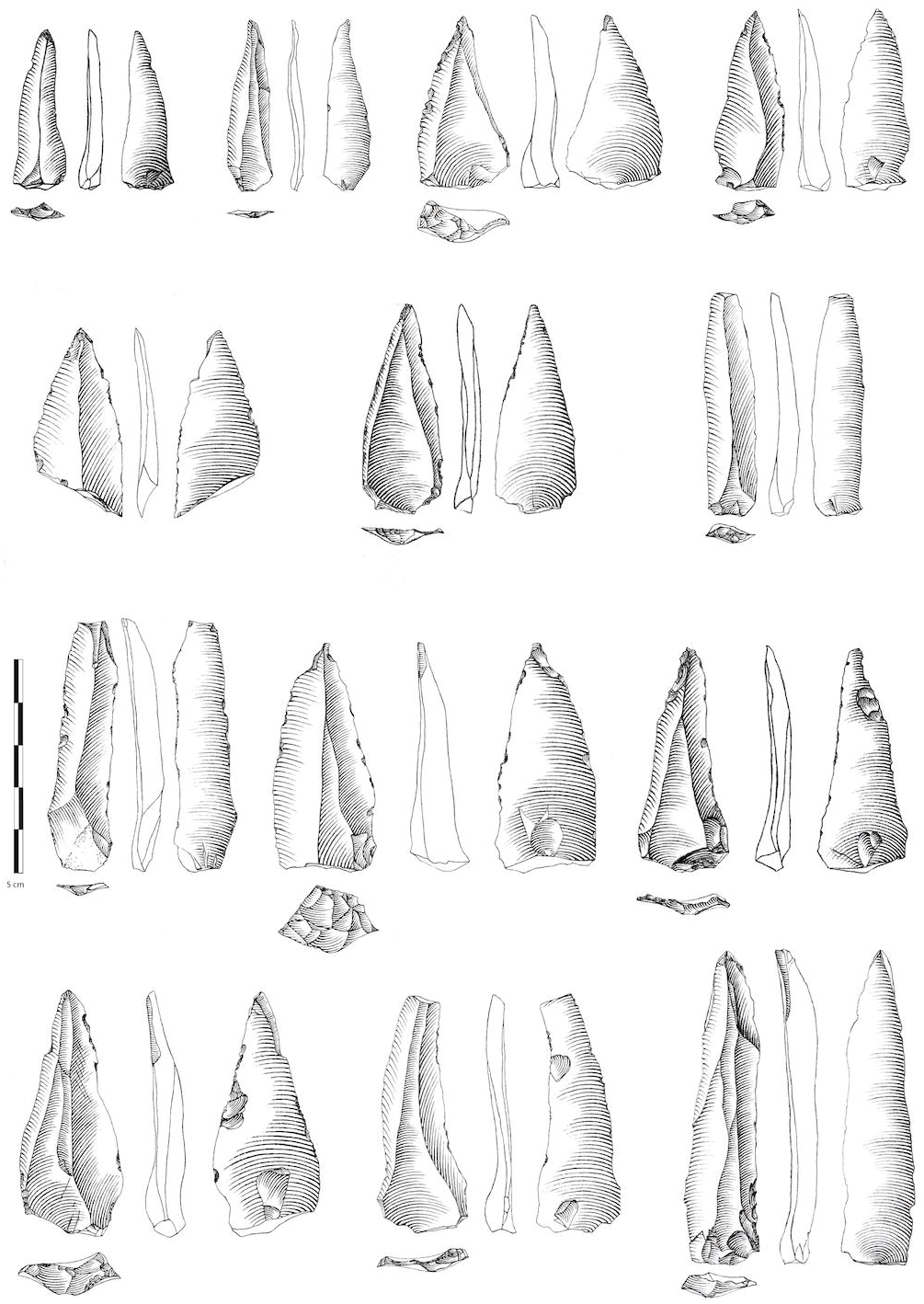
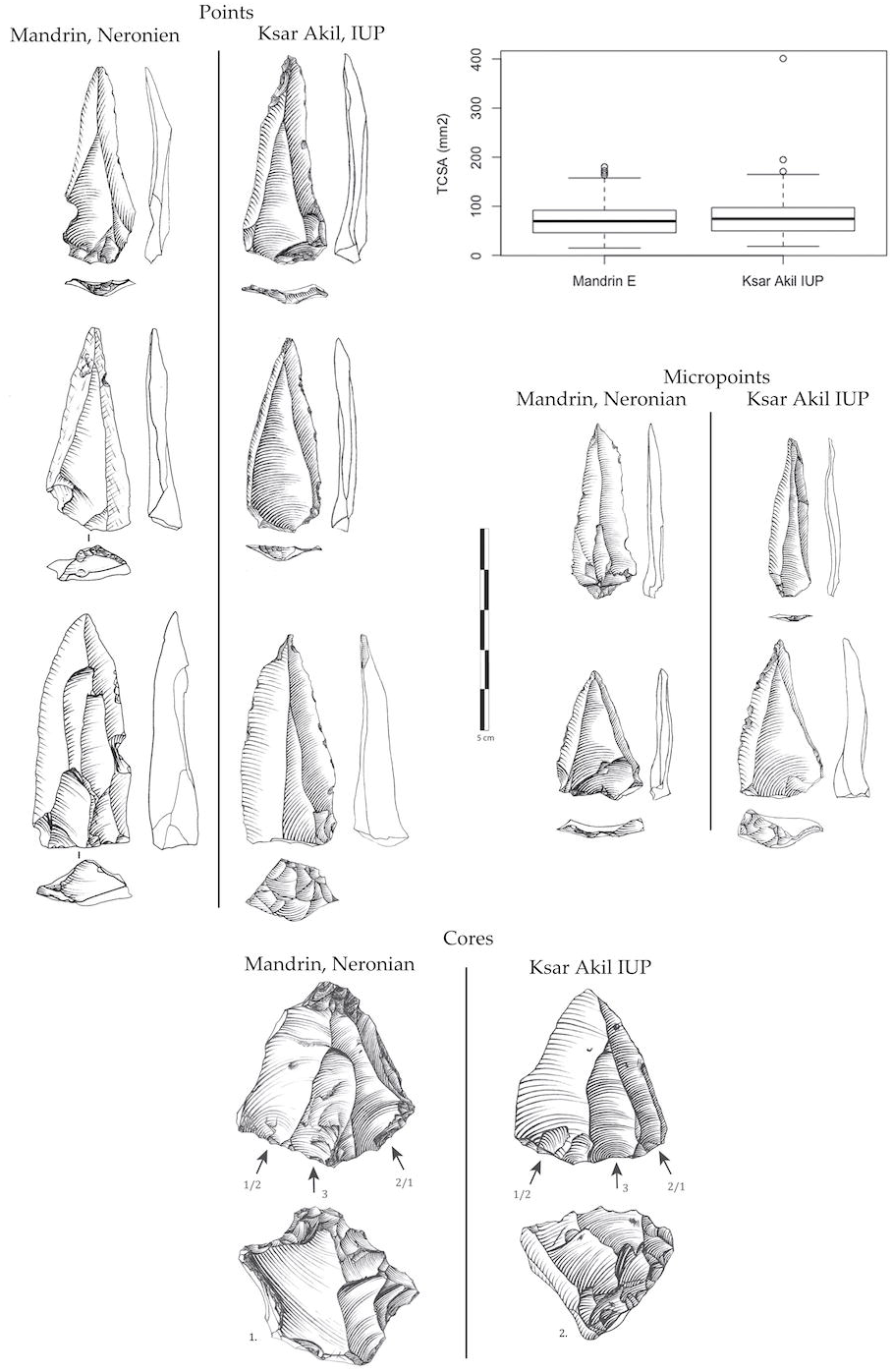
Points, micropoints, and cores from the IUP of Ksar Akil and from the Neronian of Mandrin E. The technical systems and the production objectives are strictly identical. The TCSA (width and thickness ratios) relate to measurements per mm and show no statistical difference. Drawings by L. Metz and L. Slimak.

Recorded in Western European sequences, we then have phases I and III, from layers XXV to XX, then XIII (or even XV/XIV) to XI respectively, of Ksar Akil. The two highlights of the early Levantine UP thus find a direct and very precise echo in Western Europe through the Neronian and then the Protoaurignacian. What about Ksar Akil’s second phase from layers XIX to XVI? Would there not exist, in Western Europe, an initial phase of the UP organized around the debitage of small blades obtained by essentially bipolar debitage and turned toward obtaining backed points? The debate on the origin of the Châtelperronian and its technical relations with preceding and succeeding industries began with Breuil [54] and continued throughout the 20^th^ century [20, 55-58]. Today, it opposes two schools of thought that either consider the Châtelperronian as a full UP that has no real roots in the local industries of the Mousterian [59-64], or as a local product resulting from the evolution of preceding local Mousterian [46, 65-68]. In this debate, the question of backed points occupies a central place, as does the supposed absence of backed points or pointed blades in Mousterian collections located outside the range of the Châtelperronian and other early UP complexes [68; Supplementary Note 7). On this issue, the demonstration can be considered particularly fragile since it focuses for the Mousterian on “elongated backed blanks/points,” a category which, through the approach of those authors, technically encompasses any morphologically slender support and not specifically products resulting from blade debitage *sensu stricto*. Nor do the authors typologically associate these blanks to any shape of back-backed, cortical, *débordant*-in order to balance this assembly of technically and typologically distinct characters (natural, cortical, and backed backs) with Châtelperronian points. However, Châtelperronian points are well-circumscribed blanks as to their technical and typological nature which only very partially overlap this definition. The Châtelperronian point concerns true blades-technically exclusively obtained from blade debitages- and then sharpened with various forms of abrupt retouching, or even true truncations. This point alone highlights the fact that these comparisons between Châtelperronian and local industries in the Mousterian are largely based on aspects which remain rather superficial, from the point of view of the technical systems involved, mainly based on morphological properties and not on the precise technical structures present (Supplementary Note 7). Meanwhile, true backed points represent precisely one of the structural elements of NEA technical systems. More precisely, if one is technically and typologically rigorous on the definition of the backed point therefore concerning, like in the Châtelperronian, exclusively true blades associated with backed backs, these technically well-circumscribed products do not structure, on the scale of Western Eurasia, any other industry than the NEA, which has until now been completely interpreted as one of the Levantine counterparts to the Protoaurignacian. Analysis of the Ksar Akil sets shows a significant number of points that are technically and typologically undistinguishable from those of Châtelperron. The craftsman of these backed point industries, *H. sapiens*, was found in layers XVI/XVIII of Ksar Akil (Supplementary Note 3). It is remarkable that these sets were classified as early as 1947 by Father Ewing as Châtelperronian, a classification which disappears in later studies. They are stratigraphically positioned between the IUP-technically similar to the Neronian, and the XIV-XI assemblages-technically similar to the Protoaurignacian. Identifying the hominin makers of the Châtelperronian remains uncertain as it is still reliant on data from older excavations plagued by stratigraphic uncertainties (e.g., Arcy sur Cure). *H. sapiens* are biologically recognized in the Neronian/ IUP of Mandrin E [1], in the EUP of Ksar Akil [11, 15], and at Bacho Kiro [69], and in the European Protoaurignacian [70, 71]. We can therefore highlight that the three key phases of the first UP of Ksar Akil have clear parallels in contemporary industries in Western Europe, recognized from the Rhône Valley to Franco-Cantabria.

These elements make it possible to posit an original thesis on the origin, structure, and evolution of the first moments of the UP in Europe where we would see recorded horizontally (geographically) in space, what is recorded vertically (stratigraphically) at Ksar Akil. The state of the archaeological documentation does not make it possible to link, from one person to another, Levantine and European spaces that would appear to be isolated in terms of the content of their records. This state of affairs affects the three major phases of the division that I propose of this first UP, even if ensembles such as Bacho Kiro or Temnata could be interpreted as intermediate points between East and West [31, 44]. In the model proposed here, the elements of the Bachokirian would not correspond precisely to the Levantine IUP as recently proposed [69], but rather to one of the initial phases of the EUP, and thus to the beginning of the NEA, prior to the full development of the backed points, whereas the Châtelperronian would correspond to a more evolved stratum, therefore more recent, of this same phase of the EUP (Supplementary Notes 2 & 6). The equivalent of phases I and II of Ksar Akil, which correspond to the IUP and the full development of the EUP, would thus only be currently documented at the western extremity of Europe, on the Mediterranean and Atlantic façades of France and the Iberian Peninsula. One should note the absence of data on the first UP coming from the Turkish peninsula outside its Levantine comma of Hatay, an absence that is directly incumbent on the history of research in this geographic space [72]. The significance of lack of data in this key area have long been recognized, as have the implications on Mediterranean correlations which have hitherto been limited to the Protoaurignacian and Early Aurignacian [29]. The northern Mediterranean does not document the three articulations that we see at the eastern and western ends, suggesting the existence of maritime routes linking the two sides starting at least 55 ka. Although direct evidence of long-distance maritime navigation capacities are not clearly demonstrated in the Mediterranean until after the Last Glacial Maximum [73], they are now little-questioned at the opposite eastern end of Eurasia during the peopling of Sahul starting 65 ka [74, 75].

The sequence of Ksar Akil allows us to document the precise technical emergence of industries identical to the Protoaurignacian of Europe (SEA), a development that can be broken down into three successive technical stages resulting from a progressive evolution of the technical systems of the first Levantine UP; IUP/ NEA/ SEA. These successions in the stratigraphy have remarkable parallels with the western end of Europe with the Neronian/ Châtelperronian/ Protoaurignacian triptych. Across western Europe, from France to Iberia, we would then have a technical and cultural structure identical to that recognized in stratigraphic successions, through time, in the Eastern Mediterranean. Radiometric analyses show an indisputable chronological anteriority of the Neronian over the Châtelperronian [1], and one can also therefore reasonably posit the chronological anteriority of the first phases of the Châtelperronian over the Protoaurignacian.

To resume, based on the analysis of the technical structures of the Ksar Akil sequence, I propose that the three phases of the first Levantine Upper Paleolithic find a strict corollary across Europe:

-Phase I, corresponding to the IUP, with points and blades, potentially begins in the middle of the 50th millennium and is recognized in only a few sequences in Europe, including the Neronian, the Bohunician and the Kremenician, across discontinuous spaces from the Rhône valley to Ukraine. The IUP *sensu stricto*, with points and micropoints and unipolar debitage, as documented in the Ksar Akil sequence (XXV-XXII), is only documented in Rhône area with the Neronian. A variant of this IUP, *sensu lato*, with large points and bipolar debitage is well attested in the base levels of Boker Tachtit; their links with the Bohunician have precisely been approached by Tostevin. It is not possible at this time to define whether we are confronted here with a synchronic cultural diversity or with two evolutionary stages of this IUP.

-Phase II, corresponding to EUP I / NEA, is characterized by its production of small bipolar blades and backed points. The NEA finds singular technical correspondence with Châtelperronian productions. Its geographic distribution is clearly different from Phase I and now affects the French Iberian and Atlantic areas. The Bachokirian, weakly bipolar and not characterized either by the representation of points *sensu* Levallois, nor by backed points, could correspond to the first stages of the NEA as documented at Ksar Akil (XIX-XX); it would then be slightly earlier than the Châtelperronian in the west of the continent, before full developmental phases of the backed point at Ksar Akil (XVII-XVI).

-Phase III, EUP II/SEA/Protoaurignacian, focuses on the production of long rectilinear bladelets obtained by unipolar debitage. These industries are recognizable in all regions from western Europe to the Levant, uniting for the first time all Western Eurasia.

These three phases from the beginning of the Upper Paleolithic can be interpreted as three distinct migratory waves of biologically modern populations that systematically had their origin within the Mediterranean Levant, where different sequences make it possible to document the gradual emergence of phases II and III from the local cultural substrate.

### From colonization to relations with the Neandertals, what distribution models for the first *H. sapiens* in Europe?

If in the Levantine space the emergence of the SEA/ Protoaurignacian makes it possible to recognize its emergence in three clear stages, this entity is structured based on a gradation of technical systems originally focused on obtaining slender Levallois points from unipolar debitages. Here we have indications of continuities in traditions, and probably also of biological populations in the broad sense. It does not seem possible to document such continuity in the Western European area. We do not recognize any sequences that allow us to perceive a progressive evolution from the Neronian to the Châtelperronian then to the Protoaurignacian. We do not know of any other collections that could be considered to present intermediate technical indicators between these three industries. At the same time, we note that these industries differ not only in time, with the Neronian/ Châtelperronian / Protoaurignacian successions, but also in their spatial distributions; Middle Rhône / Atlantic France-Iberia / Western Europe (Fig. 9). This spatio-temporal succession shows geographic distributions that are both disjointed and increasingly vast. And for the first time, the third phase, the Protoaurignacian, culturally unites the western European space and the Mediterranean Levant.

**Fig. 9.**
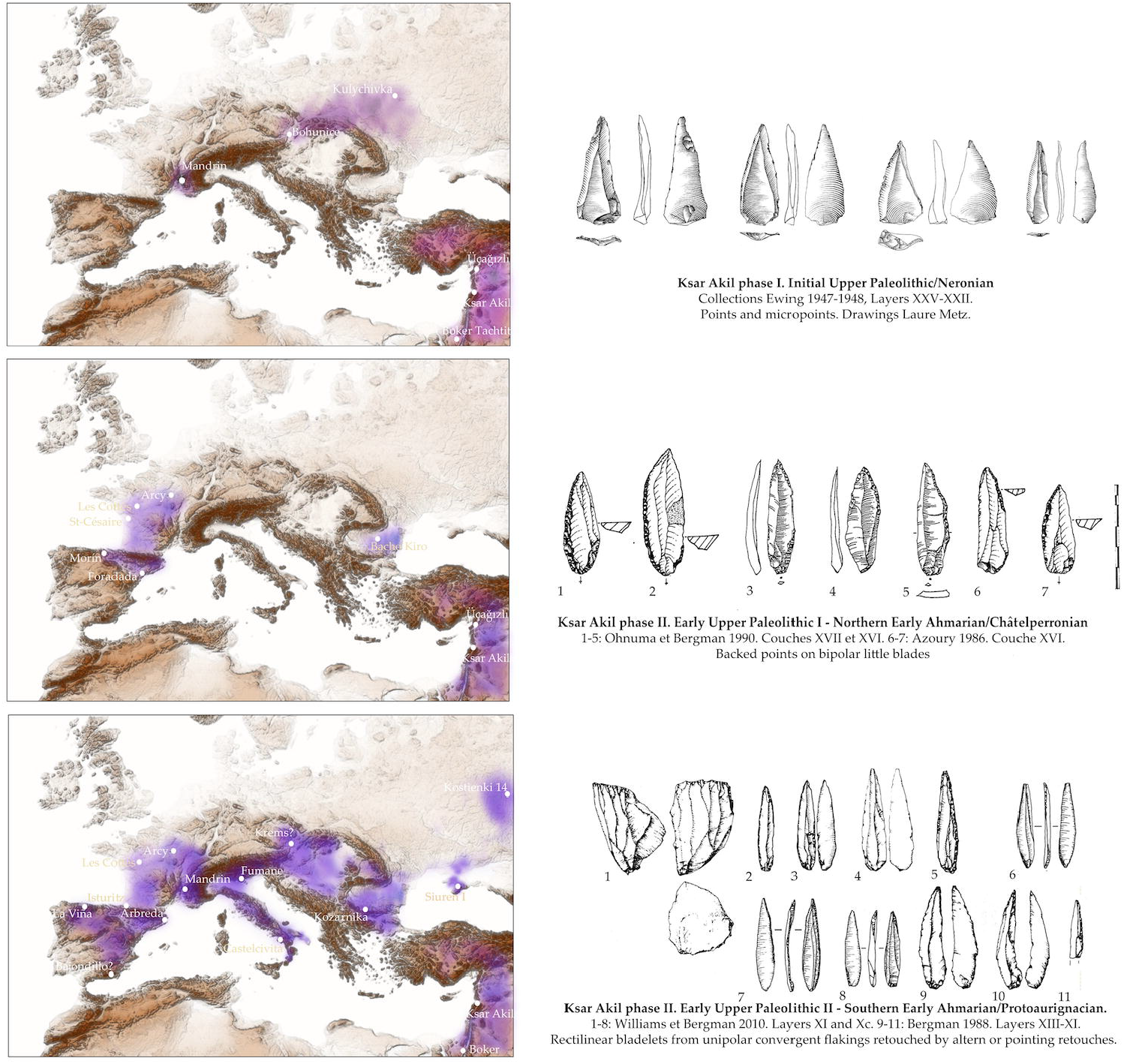
Based on the analysis of the technical structures of the Ksar Akil sequence, I propose that the three phases of the first Levantine Upper Paleolithic find strict corollaries across Europe.

Here, these temporal and spatial peculiarities have every reason to be interpreted as the archaeological signature of three distinct migratory phases, all likely stemming from the same Levantine cultural substrate. The first migratory phase is relatively old, prior to 54 ka, and is currently recognized only in the Rhône valley. It can be noted that the Rhône is the main natural artery connecting the Mediterranean area with the great steppes of northern Europe. If we start from the observation that the IUP groups are familiar with Mediterranean maritime areas, as evidenced from the distribution of Levantine sites, we can also note that this first migratory phase does not seem to move away from the Mediterranean shores for more than a hundred kilometers. At the same time, the records from Grotte Mandrin allow us to document a continuous presence of these populations for around forty years in this territory [4]; the equivalent of a human generation or two and no more. This first migratory phase apparently abandoned this territory without leaving behind discernible biological or cultural descendants. Whatever the reason for the abandonment, we can decipher important elements about this group’s structure and goals. These *H. sapiens* were probably not a simple group of scouts, but rather that their displacement had an underlying desire to permanently settle in these lands. The length of time that the group occupied this territory does not agree with a simple stop, nor with the simple desire to explore an unknown territory. The Neronian level of Mandrin yielded a tooth of a very young *H. sapiens* child [1]. This group was therefore made up of men, women, and young children, whether they were part of the trip or were conceived within these new territories. The mastery of the two banks of the river and the knowledge of all the siliceous resources over a relatively large area allow us to envisage close relations either with Neanderthal aboriginal groups, or with isolated Neanderthal individuals possessing prior knowledge of these territories [7].

The Châtelperronian would then correspond to a second migratory phase, which is only archaeologically visible several millennia later, around 45 ka. The data from Cova Foradada show that this second phase does not concern the French Atlantic area and the Cantabrian cornice alone [76] but was also expressed as far as the Iberian Mediterranean area, far away from the distribution territories of any Mousterian of Acheulean Tradition which has been sometimes considered as a local antecedent to these industries [46, 65, 67, 68]. If the Châtelperronian effectively corresponds to a second migratory phase by *H. sapiens*, and originated from the same Levantine cultural substrate, the absence of chronological and geographical overlap between phase I (IUP / Neronian) and phase II (NEA / Châtelperronian) is all the more remarkable, as the territorial expansion of this phase II affected large territories-Atlantic, continental, and Mediterranean-which remain quite geographically disjointed. Over this same period, the Rhône valley was occupied by Neandertal groups that carried the Post-Neronian II traditions [1]. Could it be that in the same geographical space that saw the first migrations of *H. sapiens* into Europe, Neanderthal groups no longer allowed access to their previous territory? This would be remarkable, since the Post-Neronian I and Post-Neronian II, which mark a return of Neandertal populations to a large territory around Mandrin, also indicate a persistence of Neandertal populations in one of the main migratory arteries of Western Europe [1]. This could well indicate a refusal or a resistance from the aboriginal populations against a return of *H. sapiens* at the very moment when, according to this hypothesis, these latter populations would manifest their first real colonization by way of settlements, not only numerous, but also over vast territories across Western Europe.

The Protoaurignacian, the third phase, nonetheless remains the first real layer of *H. sapiens* populations to be expressed over all of Europe and as far as the Mediterranean Levant, marking the cultural and territorial unification of these groups across the continent. It is only in this phase III that the native Neanderthal populations were replaced by *H. sapiens* populations. This replacement process was expressed not only over a few generations but on a Western European scale and even at very geographically specific areas, such as the Rhône valley, for over at least 12 millennia.

The comparative analysis of this trans-Mediterranean documentation suggests processes that are not only long, but also non-linear. This includes records of successive phases of contacts, and of cultural replacements in well-defined territories. In the Rhône Valley, these contact / replacement / exclusion processes are expressed precisely in 4 biological stages and 5 cultural stages (from oldest to youngest); Rhodanian Quina (Neanderthals) / Neronian (*H. sapiens*) / Post-Neronian I (Neanderthals) / Post-Neronian II (Neanderthals) / Protoaurignacian (*H. sapiens*) [1]. We know from the soot analysis [4] that in the two instances where *H. sapiens* are present at Grotte Mandrin, the time between the Neanderthal installations and the *H. sapiens*’ is only a few seasons, possibly only one year. In the phases directly preceding the *H. sapiens* installations, by seeing the extent of the perceptible territories of these groups and of the recurrences of seasonal Neanderthal installations in the cavity over several decades, we can here parsimoniously ask if, in this very particular place of the middle Rhône valley, in this very cave, or in its immediate surroundings, these unique archaeological records imply the existence of direct contact between populations. With the probability that the migrant populations benefited from the knowledge of the aboriginal populations, we can perceive the most direct implications within the *H. sapiens* groups, allowing a precise knowledge of the resources from this rather vast territory [1]. The precise nature of these transmission processes from Neanderthals to *H. sapiens* is not directly perceptible. The possibility of Neanderthal guides integrated within the *H. sapiens* group could be seen as both minimal interpretation and universally documented in ethnography.

There remains the enigma of the middle valley of the Rhône, not only having recorded the first migration of *H. sapiens* into Europe, 10 to 12 millennia prior to the first migrations hitherto recognized, but also the only occurrence of successive biological replacements in Eurasia-Neanderthal / *H. sapiens* / Neanderthal / *H. sapiens*. The structure of these replacements and the first arrival of *H. sapiens* in the heart of mainland Europe’s main north-south river artery can hardly be considered anecdotal. The similarity with the Levantine productions suggests the existence of maritime movement networks which would have already been solidly in place from 54 ka. The absence of the second migratory phase, NEA/Châtelperronian, from the Rhône area and being framing by the southwest, the west and the north would be equally remarkable, the territories that were occupied again by Neanderthal populations were no longer appearing accessible to the *H. sapiens* populations.

This pattern of colonization in Europe and replacement of local populations accounts for an important part of the cultural facts recorded in Western Europe during this 10 ka period. Along with the Uluzzian in the western Mediterranean [70], the presence of *H. sapiens* groups with clearly distinct traditions underlines the cultural richness of these population replacement processes. This would also indicate that, whatever the relations between *H. sapiens* and Neanderthals in phase II, as soon as 45 ka Western Europe would already have been largely occupied by different *H. sapiens* populations. At the same time, these data indicate that the last Neanderthal populations do not appear to retreat into refuge spaces, but actually occupy without sharing for a few millennia, major axes of circulation on the scale of the European continent. The technical structures of these societies do not allow us to document any obvious form of acculturation from *H. sapiens* to Neanderthals, except perhaps the precise knowledge of certain technical know-how related to point production technologies [7], which could well have been acquired by the aboriginal groups during the very first migratory phase of *H. sapiens*. The last Neanderthal populations would then not only be bearers of their technical traditions, relatively immutable over hundreds of millennia, but would also, paradoxically, be the only heirs of technical traditions long abandoned by *H. sapiens* and belonging to the first phases of settlement in Europe by these populations. It is in this light of equivocal conservatism that these final Neanderthal populations will definitively take their ‘révérence’, replaced in just a few seasons, as indicated by the soot records, by a wave of a population which will finally unite Europe, reaching a tipping point into the historical structures of the UP.

## Acknowledgments

Long-term research at Grotte Mandrin was made possible support from of the Regional Archaeological Services in Lyon and many locals of Malataverne (Drôme). Analysis of Ksar Akil’s collections was made possible by the staff of the Peabody Museum in Cambridge, Massachusetts (especially Jeffrey Quilter, Viva Fisher, Kara Schneiderman, Laura Costello, Diana Loren, Lainie Schultz, Emily Pierce Rose, Meredith Vasta, and Diana Zlatanovski), Christian Tryon, and support from the American School for Prehistoric Research to organize the collections, and logistical support from the InSHS of the CNRS. A Harvard University Radcliffe Institute seminar organized by Laure Metz and Christian Tryon in 2019 made it possible to present the first frameworks of these relations between East and West at the beginning of the UP. Numerous exchanges with Christopher A. Bergman were crucial to understand the history of research at Ksar Akil. Jason Lewis and Christian Tryon provided invaluable editorial support.

## Supporting Information

S1 File. Contains:

Supplementary Note 1. Evolving Thoughts on the Origins of the Neronian Supplementary Note 2. History of Correlations between European & Levantine

Archaeological Sequences

Supplementary Note 3. Radiometric Dating of the Ksar Akil Sequence

Supplementary Note 4: Salient features of the Technical Structures of the IUP and EUP at Ksar Akil

Supplementary Note 5: After the EUP of Ksar Akil

Supplementary Note 6: From East and West. Back to Mandrin, downgrading, reclassification, pieces of the puzzle of Western Europe

Supplementary Note 7: The Châtelperronian Question

Supplementary References

