## Supplementary Text for "The Three Waves: Rethinking the Structure of the first Upper Paleolithic in Western Eurasia"

Ludovic Slimak

Corresponding author = Ludovic Slimak

**This PDF file includes:**

Supplementary Text:

Supplementary Note 1. Evolving Thoughts on the Origins of the Neronian

Supplementary Note 2. History of Correlations between European & Levantine Archaeological Sequences

Supplementary Note 3. Radiometric Dating of the Ksar Akil Sequence

Supplementary Note 4: Salient features of the Technical Structures of the IUP and EUP at Ksar Akil

Supplementary Note 5: After the EUP of Ksar Akil

Supplementary Note 6: From East and West. Back to Mandrin, downgrading, reclassification, pieces of the puzzle of Western Europe

Supplementary Note 7: The Châtelperronian Question

Supplementary References

**Supporting Information**

### Supplementary Note 1: Evolving Thoughts on the Origins of the Neronian

The hypothesis put forward in 2004 [7] of a local origin of the Neronian and of a continuity between the Rhodanian Quina and the Neronian was based mainly on data from the 1950s excavations at Grotte de Neron by Jean Combier and Maurice Veyrier. In fact, in the Rhodanian Quina collections of Grotte de Neron one finds blades and points that are technologically identical to those in the overlaying Neronian level. At Mandrin, the Rhodanian Quina has the same position in the stratigraphy, in level F under the Neronian, and of remarkable stratigraphic integrity. The technological structures of the Rhodanian Quina of Neron and Mandrin are identical [1], reinforcing the cultural value of these regional facies of the Mousterian considered 23 years ago [80]. Contrary to the Quina from Neron layer II, the data from Mandrin F does not reveal any piece that is technologically identical to the rich assemblages of blades and points that overlie them. While waiting to receive a richer corpus on this Rhodanian Quina through additional excavations elsewhere, it therefore appears reasonable to abandon for now the hypothesis of continuity between this local Mousterian and the Neronian [1]. Meanwhile, comparative analysis of Mandrin E and Ksar Akil's IUP levels shows a remarkable technological identity. It is based on these two lines of information that it was initially proposed in 2017 and 2019 that these Neronian industries could represent a *H. sapiens* incursion of Levantine origin into Neandertal territories [5, 6]. It should be noted that these hypotheses were published one and a half years before hominin teeth from various levels at Mandrin were analyzed and taxonomically attributed. These conclusions from cultural and biological anthropology both point today to a *H. sapiens* origin for the Neronian- conclusions that were drawn from separate and distinct analyses that both independently led to identical conclusions. The reversal of my own hypotheses is therefore based exclusively on the deepening of technological knowledge concerning the different assemblages from the end of the Middle Paleolithic and from the beginning of the Upper Paleolithic and the increased stratigraphic resolution provided by the ongoing excavations at Grotte Mandrin relative to those done in the 1950s at the Grotte de Neron. It is, in no case, a re-articulation of my own thoughts according to the results of paleoanthropological analyses. This point has significant historiographic value all the while highlighting the predictive nature of analyzing technological systems, which pointed to a *H. sapiens* origin prior to the analysis of human remains from the Grotte Mandrin sequence. The data on the origin of the Neronian therefore seems to be absent from the western Mediterranean basin, thus inducing unique anthropological, biological, and cultural questions that must be considered.

**Supplementary Note 2: History of Correlations between European & Levantine Archaeological Sequences**

Following a general statement made by Ofer Bar-Yosef on the relations between the Protoaurignacian and the Levantine Ahmarian [81], Mellars first points out that "the most convincing origins for these technologies seem to occur in sites in the Near East (for example, in the lower levels of the Ksar Akil sequence in Lebanon, or at a number of open-air sites such as Boker A" [28], a wording reused in 2006 by Zilhão who proposed that "Technologically and typologically, the Protoaurignacian is virtually indistinguishable from the Early Ahmarian of the Levant" [32]. Here, Zilhão precisely refers to the industries of Kebara, Ksar Akil, and Üçagızlı, that is to say the industries of the North Levantine area (Northern Early Ahmarian, NEA), for which Ksar Akil represents the type sequence [ref. 32, p. 188, table 1, and p. 190]. Since then, these comparisons have been systematically repeated *in extenso* in studies of the beginning of the Upper Paleolithic [e.g. refs. 42-46][*], but have never been supported by a direct analysis of the two

[*]"Similarities between Proto-Aurignacian and Early Ahmarian assemblages are particularly significant in terms of blade and bladelet core reduction methods and retouched bladelet morphologies (e.g. certain El-Wad points resemble the Font-Yves points of the Proto-Aurignacian, Belfer-Cohen, Goring-Morris 2003). The convergences are also of particular significance when examining the general "allure" of blade and bladelet blanks, often standardized and regular, narrow and elongated and with a predominant rectilinear profile. All these technological and stylistic patterns well differentiate the Early Ahmarian and the Proto-Aurignacian on the one hand from the classical Early Aurignacian on the other hand. Moreover, as in the Proto-Aurignacian, the Early Ahmarian industries include few examples of organic productions and the predominant use of shells for ornaments, as recently demonstrated in **levels F–H of Üçagizli** for instance (Kuhn et al. 2003)" (Teyssandier 2006). Here there is an association of the Protoaurignacian with the SEA in first part of the citation, and with the IUP of Üçagızlı in the last sentence, bolding ours.
"They –Emiran and IUP- appear, however, to represent part of a local developmental continuum that subsequently yielded a related industry (Ahmarian) associated with modern human remains in **layer XVII at Ksar Akil** (…). The Proto-Aurignacian seems to have close ties with the Ahmarian industry of the Near East in much the same way that the Bohunician is tied to the preceding Emiran. As already noted, the Ahmarian is associated with modern human remains" (Hoffecker 2009).
"Les schémas opératoires identifiés, leur association à ces deux types de lamelles retouchées, la présence de parure (coquillage perforé, dents percées) ainsi que la position chronostratigraphique de l'industrie plaide en faveur d'une certaine correspondance avec les premières industries du Paléolithique supérieur, centrées autour d'une intention

industry groups. These comparisons are therefore grounded on bibliographical reference to statements about the technical traits that are supposed to characterize the Protoaurignacian and the Early Ahmarian, the notion of the Early Ahmarian systematically encompassing the sets of the NEA from the sequences of Ksar Akil and Üçağızlı.

Yet, in two more in-depth studies, but still without direct ties to the collections, Mellars proposed a comparison of the Protoaurignacian with notably more recent sets from Ksar Akil, which he attributed to the “Levantine Aurignacian B”, in units located more than 2 m stratigraphically above the sets that Zilhão referred to; “*These bladelet types are particularly frequent in the so-called “Levantine Aurignacian B” assemblages from levels 9-11 at Ksar Akil”* [29]. Mellars also referred to the industries of Boker A from the Negev Desert in Southern Israel, and thus to the industries of the Southern Early Ahmarian- SEA [82, 83].

A Levantine origin for the Protoaurignacian lamellar sets was commonly highlighted by Mellars in 2004 and Zilhão in 2006. But Zilhão compares units XVI to XX of Ksar Akil, while Mellars compares with elements of composite origin, thereby grouping the industries of Boker A of the SEA and units XI to IX of Ksar Akil, and wrongly attributed in his study of “Levantine Aurignacian B” [29]. In defense of these authors, and of those who followed them over the subsequent years, insisting on the historiographical richness of the research from Ksar Akil, whose simplified synthetic overview is provided in Fig. 1, permits us to perceive its complexity. This figure focuses

lamellaire et lamino-lamellaire forte. Tel est le cas, par exemple, dans le Proto-Aurignacien d’Europe occidentale, l’Ahmarien ancien du Levant ou le Baradostien du Zagros.(…) L’Ahmarien ancien est connu du nord au sud du Levant (…) Son industrie se caractérise par une production normalisée de lames et de lamelles, débitées par percussion directe tangentielle au percuteur tendre. Des variabilités régionales se dégagent des ensembles lithiques, montrant l’existence probable de deux faciès géographiques (…) l’Ahmarien septentrional apparaît à **Üçagizli B,B1-4,C, Ksar Akil XX-XV**” (Tsanova et al. 2013). There is a recognition of the two facies of the Early Ahmarian, but they essentially confuse the technical characteristics of the SEA, the industries of the NEA are also seen to be characterized by “un débitage unipolaire convergent, prismatique ou pyramidal, en vue d’obtenir des supports lamino-lamellaires élancés et pointus, de profil rectiligne à légèrement incurve” (Tsanova et al. 2013 see table 4 from the same article).

“If one assumes that the **Early Ahmarian/Kozarnikian/Protoaurignacian** represent a consistent wave of peoplings in western Eurasia, the identity of the makers of this assemblage would be primarily substantiated by the discovery of an immature individual in **layer XVII of Ksar Akil** (Lebanon)” (Hublin 2015) and see fig.3 where NEA and SEA are considered as a whole and associated with the Protoaurignacian.

“Dufour bladelets are abundant in all Proto-Aurignacian sites, including the earliest ones (e.g., Fumane: [Broglio et al., 2005]). Similar retouched bladelets are well represented in the early Ahmarian of the Levant, an industry which might be connected anthropologically and chronologically to the Proto-Aurignacian (Hoffecker, 2009)” (Roussel et al. 2016). Here, the correlation is established via the data from Hoffecker, therefore encompassing the Northern Early Ahmarian of Ksar Akil XVII in the industries associates with the Protoaurignacian in Europe.

on the main conclusions of the authors who have directly examined the Ksar Akil collections, and builds on similar work by Bergman [e.g. refs. 37, 79]. Over 70 years, from 1947 to 2017, one can see the evolution of the different cultural phases attributed, revealing significant fluctuations both in the appellations and in the boundaries of the different phases. Later I will come back to what a classic “French” technological reading can bring to the understanding of these technical and cultural successions.

After 2003 the Early Ahmarian/Protoaurignacian correlation summarizes, in the bibliography, the technical connections between East and West with episodic inclusion in the eastern group of a few other SW Asian industries such as the Baradostian [84-87] or Iranian Rostamian [88-90]. It is on this simplified correlation of Protoaurignacian = Early Ahmarian that the writings from the past 17 years have rested, generally not establishing any distinction between Southern Early Ahmarian (SEA) and Northern Early Ahmarian (NEA), but often systematically referring to the collections of the Early Ahmarian of Ksar Akil, and so the NEA. These correlations with the Early Ahmarian, which emerged as soon as 2004, were nevertheless established without directly returning to the collections, but by relying, according to the authors, on clearly distinct stratigraphic and technical facts comprised between units XX and IX of Ksar Akil, which are, in fact, expressed over a stratigraphic thickness of more than 5 m. The stratigraphic inaccuracies and hesitations vis-à-vis the key sequence of Ksar Akil therefore had repercussions on most of the subsequent studies which aimed to define the chronological antecedence of either the Early Ahmarian from the Levant or the Protoaurignacian from Europe. These studies focused mainly on the chronology of the Protoaurignacian, in part compared to levels XIV-XXV of Ksar Akil, encompassing the Initial Upper Paleolithic from levels XXV-XXI and the Early Upper Paleolithic (Early Ahmarian) from levels XX-XIV [14, 50, 51, 91].

The distinction between the NEA and SEA, whose technological divergences have been understood for some time [92] are not reflected in either radiometric studies nor in technical comparisons. The Protoaurignacian finds itself systematically associated with a syncretic notion of the Early Ahmarian which, as described above, encompasses the SEA and the NEA as encountered at Ksar Akil [42-46]. It should be noted that the deep technical differences between NEA and SEA were underlined in 1980 during the colloquium “Prehistory of the Levant” in Lyon in which Jacques Tixier also participated [18]. Jim Phillips, who was working on the industries of the Early Ahmarian in the Abu Noshra basin in the Sinai region in the South Levant and analyzing the Ahmarian XVII and XVI levels of Ksar Akil, exclaimed “my material doesn’t look anything like this” [93]. The technological distinctions between the two industries, which are indeed very marked, were also reaffirmed by all the participants of the “Prehistory of the Levant II” colloquium, held at the Maison de l’Orient-Lyon in 1988 [93].

Since then, the NEA and SEA have always been clearly differentiated based on their technological and typological structures [94-96]. We will also note that some studies [37, 47] allow us to state parsimoniously that NEA and SEA may not correspond to geographic variations of the same culture group but to chronological distinctions, with the SEA related to more recent phases of Ksar Akil as recorded in X-IX levels [ref. 37, phase 4; Fig. 1].

The Bachokirian industries have been brought closer to the Levantine IUP [31, 69]. However, these industries do not possess any of the structural features of the Levantine IUP, the most salient technical feature recognizable in Ksar Akil’s ensembles being the systematic production of Levallois-type points resulting from convergent unipolar flaking. These points, at the heart of the IUP systems of Ksar Akil and Üçağızlı, are strictly absent from these Bachokirian series. The Bachokirian is an industry with blades resulting from parallel, non-convergent flakings, and described as marked by “the frequency of bipolar operations conducted from two opposite striking platforms. Close to 20% of laminar blanks with parallel anterior negatives bear witness to this: they carry at least one anterior laminar negative of opposite direction” [97]. In recent excavations, Layer I again delivered no Levallois-type points, and shows a blade industry resulting from non-convergent parallel debitage [69]. The bipolar fraction is less clearly represented in this small corpus but remains visible on 3% of these blanks. However, the bipolarity of this debitage is probably more important. “The blades of bipolar debitage models are half as many (n=111) than the unipolar blades. We suppose that bipolar removal pattern debitage is assumed to have lower visibility due to their high fragmentation” [31]. At the same time, the absence of any Levallois-type point excludes any direct connection with IUP sensu stricto, whereas the bipolar fraction of this debitage directs us toward the initial phases of EUP/NEA as documented in the Ksar Akil sequence. The same is true for the strong representation of the retouched points at the heart of the tool corpus that would bring this industry closer to Phase II, EUP/NEA, of Ksar Akil.

**Supplementary Note 3: Radiometric Dating of the Ksar Akil Sequence**

The chronologically stricter studies mainly oriented toward the key sequence of Ksar Akil found themselves confronted with severe interpretative dead-ends stemming from divergent radiometric results [14, 50, 51, 91] and attempts to correlate Europe and the Levant by comparing either the cultural phases (Early Ahmarian from Ksar Akil *vs* Protoaurignacian; refs. 14, 98) or the biological-cultural attributions (*H. sapiens* Egbert - Ksar Akil layer XVI- *vs* the Aurignacian in Europe; ref. 50)[†], but without the direct characterization of precise technical structures from these different units. In both cases, these studies essentially come back to comparing the chronological positions of the Protoaurignacian and the Aurignacian in Europe with the elements of the EUP of Ksar Akil (so the NEA) levels XVI to XX, then following the correlations unanimously employed after Mellars in 2004. However, direct analysis of the structure of these industries shows that these comparisons do not relate to technically comparable sets, and that these approaches can only lead to erroneous conclusions (*infra*).

In parallel with these recent radiometric studies and the debates that followed, it appears more parsimonious to consider that in the current state of chronological analysis capacities, one cannot detangle these issues on the basis of radiometric measurements, which in the Levantine region

[†] “In most of Ewing's publications the only information provided regarding the stratigraphic location of the human bones is that they came from 11.46 m below datum; an examination of the stratigraphic section shows that this is very close to the boundary between levels XVI and XVII. However, the depth of 11.46 m refers to the base of the stone heap under which Egbert was found and Ewing notes that "most of the skeletal remains lie somewhat deeper than this". Newcomer remarks that 11.46 m below datum is deeper than the maximum depth of 11.25 m given for the stone artifacts recovered from level XVI and he concludes that "thus the burial would appear to be in level XVII or XVIII” (Bergman et al. 1989).

appear highly volatile between 35 - 55 ka. The bone collagen is systematically poorly preserved, which does not allow for robust dating, commonly forcing $^{14}C$ analysts to measure shell carbonates and charcoals [14, 50]. Whatever the reasons for these radiometric shifts [51, 91], we observe that they obtained significantly divergent measurements which mar the establishment of robust chronological models within this sequence and maybe more generally this chronological range and in this geographical space. The construction of models must therefore be based primarily on direct technological approaches to sequences recording precisely, in stratigraphy, all of the recognized technical and cultural successions. In this process of contextualization, the sequence of Ksar Akil represents a unique reference, making it possible to lay the groundwork on which the industries at the turn of the Upper Paleolithic of Western Eurasia should be able to position themselves.

**Supplementary Note 4: Salient features of the Technical Structures of the IUP and EUP at Ksar Akil**

Since 1997, the Ksar Akil IUP has been considered as deriving from units XXV and XXI. When exploring the historiography of the research on Ksar Akil, units XX and XIX were also attributed to the IUP by Father Ewing in 1947 and then by Azoury and Hodson in 1973 [99]. A rich body of literature on the IUP and EUP industries, including numerous illustrations, can be found in the syntheses by Ingrid Azoury in 1986 [100] or Katsuhiro Ohnuma in 1988 [101]. Importantly, these previous syntheses of the IUP/EUP are based almost entirely on the 1937-1938 excavations presently housed in the UK. In Fig. 1, we can see that from 1947 to 2017 these units were generally subdivided between two and four phases. The bottom of this series is now attached to the Initial Upper Paleolithic, the middle to the Early Upper Paleolithic (Early Ahmarian), and the upper section, layers XV to XIV, or XIII, depending on the authors, not being determined in terms of the technical and cultural phasing. The passage between IUP and EUP is positioned at the level of layers XVIII, XIX or XX according to the different approaches. From the 1970s, Azoury's work, and whatever the appellations chosen or their subdivisions into cultural sub-phases (B1, IIA…), the studies remain clearly convergent in their distinction between two well-identified major phases and unanimously attributed to the IUP followed by its gradual replacement by an EUP/Early Ahmarian. These marginal variations reflect the importance that each author attributes to certain technical or typological characteristics deemed diagnostic or relevant. The overall picture nevertheless appears to be quite convergent.

In the 1947-1948 Harvard collections, I counted 4231 lithic pieces constituting units XXV to XXI. These industries are essentially made up of laminar production, blades and bladelets, corresponding to the first phases in the process of obtaining Levallois-type points (Fig. 7). These units include 677 points and micropoints (*sensu* Levallois and generally unretouched) versus 1226 blades and bladelets. The flaking is initiated by a single or double sloped crest (84 blades and ridge bladelets of different categories are attested for), the extraction of which is followed by a laminar phase within which the extraction of typologically Levallois points fits. These removals are strictly unidirectional and convergent. No points with opposite negatives are recorded in these units. The geometry of the cores is commonly pyramidal, linked to a semi-rotating exploitation of the blocks. The extraction of technically highly invested points is inserted between these laminar phases, ensuring the classic predetermined character of Levallois debitage, but here associated with cores whose geometry and volumes do not in any way coincide with the classic definition of Levallois debitage [8]. Only the notion of predetermination then reminiscences in the sets, or rather a form of technical continuity, the classic productions of the Middle Paleolithic.

Small-sized points and micro-points *stricto sensu* are well represented in all these units (Fig. 4), representing 23-40% of this corpus of points. Their maximum dimensions are on average around 30 mm. The analysis of the last removal from the micropoint cores shows at the same time that a large fraction of the smallest points- between less than 30 mm and 10 mm- are found only too scarcely in the collections compared to their real place in these technical systems. At the same time, the rarity of microflakes means that it is the excavation methods that must be linked to the lack of these smallest parts. However, the production of points with a maximum length of less than 30 mm remains well documented with regards to the cores present. The techniques are based exclusively on direct hard stone percussion. Most of these points have a finely faceted butt. The points are rarely retouched and, in general, the typological tools occupy a relatively discrete place in these units.

Layers XX through XVI or, depending on the authors, XX through XIV are assigned to the EUP/ Early Ahmarian. The differences in interpretation for the last strata attributable to the EUP may result from problems of correlation between the 1937-1938 and 1947-1948 collections since units XIV to XV, perhaps even XIII to XV, collections from the 1930's are, in certain earlier summaries, considered to register a hiatus in terms of industries or occupations of the shelter (Fig. 1). The Harvard collections are comprised of 1221 lithic artifacts for units XII to XV, no hiatus can therefore be documented here, layers XII-XV yielding 12% of all the lithic elements of the EUP-XX to XIV; n= 10175.

A continuity between IUP and EUP at Ksar Akil has long been emphasized [100-102]. Within these technical systems, no rupture can effectively be documented from one layer to the next. By going up the stratigraphic units, we can clearly see that the unipolar debitages of the IUP give way, very gradually, to bipolar (bidirectional) debitages (Fig. 5). In the underlying units of the IUP, negatives of bipolar slides and coverslips are absent or marginal, their proportions being contained between 0 and 4 to 6%. These rare indications of bipolarity within the IUP also affect, but only very locally, the blanks, in part distal, corresponding then to both limited and very episodic technical gestion of the convexities, which cannot be considered as evidence of bipolar debitage in the strict sense. Figure 5 shows that in the sequence of the EUP, from units XX to XV, bipolar debitage develops very gradually to concern exactly 50% of the blanks of layer XV before falling sharply to 15.4% in layer XIV, the last phase of the EUP. This transition to bipolar debitage is accompanied by a progressive overturning of percussion techniques, direct mineral hard hammer percussions giving way to soft hammerstone, then to the progressive emergence of organic soft percussions *stricto sensu* in the upper strata of the EUP. At the same time, the faceted butt of the IUP give way to the much thinner abraded, smooth butts. Quantified approaches to these processes are available in the aforementioned summary [100, 101]. This overturning of debitage and butts encountered on the blanks between the IUP and the EUP has also been underlined and quantified within the Üçağızlı sequence in the Turkish Hatay [102]. This progressive development of soft and organic percussion induces obtaining blanks, and in particular of sharp blades/ slender points, very slightly curved, curvature which is not found in the points of the IUP, obtained by direct hard percussion. This overturning of percussion techniques also leads to a more marked slenderness in the blanks obtained. The points, *sensu* Levallois, remain typologically well attested in this Early Ahmarian, but their appearance now refers more directly to laminar forms. If I count in these units 842 points and micropoints for 3262 blades and bladelets, the name point now only refers to the identification of specific discontinuous debitage rhythms [8], making it possible to obtain pointed blades, technically predetermined, with 3 sides, and with perfect axial and transversal symmetry. This gradual morphological evolution becomes clearly marked from units XVII-XVI. The slender points obtained in these soft hammerstone and organic percussion systems could be fully classified into pointed, regular blades and bladelets if we do not specifically analyze the organization of these debitage rhythms via a reading of the visible negatives on the ventral face of the blanks and cores in particular. The gradual morphological changes from one stratum to another also underline these processes of continuity and the technical rooting of these laminar productions in well-defined systems dedicated primarily to obtaining slender points that are technically highly invested. The very notion of a point, technically linked to a notion of predetermination based on the establishment of debitage rhythms originating in Levallois spheres [8], can no longer be recognized within a technical analysis that is too generic or which would not be based on the structure of the evolution of this point debitage, diachronically, strata after strata. Therefore, the technical analysis of these stratigraphic successions shows that these objects gradually tend toward a sharp blade, all the while retaining structural elements of the organization of this debitage, recognizable from the early phases of the IUP.

Once could imagine that this graduality could be the result of excavation methods and the physical mixing of elements from different units, thus inducing a progressive balance of technical indicators. For example, the analysis of layer XXV, a very poor unit made up of only 33 lithic pieces in the collections at the Peabody Museum, marks the beginning of the IUP, effectively allowing us to propose a mixing of elements, probably between the underlying layer XXVI of the MP and the overlaying layer XXIV of the IUP. More than a third of the pieces in this unit (n=12/33) fit into systems for producing flakes, Levallois and Discoid, characteristic of the underlying MP levels. Assuming that this unit was originally sterile from a lithics perspective, the 33 pieces of unit XXV would then represent a construction from elements of units XXVI and XXIV, stratigraphically framing it. The units XXVI to XXIV, if combined, contain 1008 lithic pieces. We could therefore calculate the maximum pollution rate of layer XXV by framing units (33/ 1008x100), i.e. 3.3%. This theoretical indicator would represent here, by definition, a maximum index of mixing since we cannot exclude that these 33 parts could belong to this unit. This 3.3% mixing indicator therefore represents, in the context of unit XXV, a maximum pollution rate, an indicator which seems quite representative for the other units in this sequence. If we wish to understand the main technical features of these stratigraphic successions, a possible pollution of 3.3% from the above and underlying layers cannot obscure the major technical trends that can be highlighted for each. Several other indicators can be put in place to assess these possible inter-layer pollutions, such as the representation of Levallois flakes across the different levels of the IUP and the EUP. The Levallois flakes are indeed strongly attested in the last units of the Middle Paleolithic and almost absent from the post-Mousterian sequence. At the heart of Father Ewing's collection, the units presenting the most Levallois flakes do not exceed 6% for layer XXV, 4.2% in layer XV, and 2.5% in layer XIII, Levallois flakes being virtually absent from other units from the first UP levels (Fig. 2). Here, we obtain a "maximum mixing index". Indeed, in the context of these Levantine sequences, clear technical continuities from the Middle Paleolithic to the Upper Paleolithic have often been proposed [*e.g.* refs. 103-106, *contra e.g.* 35, 36, 107-109]. It is therefore conceivable that certain traditional Levallois flake productions persisted in the technical systems of some of these groups of the IUP and of the Levantine EUP. Starting from the postulate that these categories of supports do not belong to these units (problems during excavation, post-excavation mixing during the management of artifacts, cleaning, classifications, etc.), the maximum mixing indicator would nevertheless remain, in the worst possible configuration, strictly marginal. At the same time, I inserted in Figure 2 the same index concerning the last two Mousterian units, XXVII and XXVI, showing that in the sets preceding the IUP, the Levallois productions concern more than 25% of this lithic corpus. These technical indicators then mark a clear distinction between the components of the Mousterian (XXVII-XXVI) and the beginning of the IUP (XXV-XXIV). A mirror analysis of this indicator can be established by analyzing the representation of blades, bladelets, and points in the entire sequence from the MP to the EUP, units XXXVI to XIII (Fig. 2).

The EUP of the layers XX to XIV of Ksar Akil then corresponds to a gradual evolution of the industries from lower layers, XXV to XXI of the IUP. This evolution sees, in parallel, a gradual shift from unipolar debitage to bipolar debitage and the changes affecting percussion techniques. Here we can propose a corollary between technical (percussion techniques) and technological (the organization of the technical system) evolutions, evolutions which nevertheless, in diachrony, preserve a common technical objective; obtain slender points and sharp, very regular straight blades. The transition to soft hammer percussion indeed makes it possible to more easily obtain slender and regular blanks than debitage with hard hammer which require the repetition of perfectly calibrated movements with risk of hinges and plunging fractures. Soft hammerstone and organic percussion have a much greater tolerance, requiring less precision when applying force. These changes in percussion techniques represent then a significant relaxation of the system, particularly affecting the strategic moment of point extraction. In fact, this development therefore represents a simplification of the system in terms of the know-how necessary for its implementation. This change in techniques nevertheless induces, in return, a more pronounced curvature of the obtained blanks. The transition to truly bipolar debitage under certain circumstances makes it possible to partially counteract this slight curvature of the obtained blanks. If one balances the diachronic structure of these technical systems, which gradually evolve from unipolar to bipolar, with this emergence of new percussion techniques, the transition from IUP to EUP would ultimately only correspond to an evolution of percussion techniques in favor of soft hammerstone percussions, the entire system adapting to the constraints induced by these new percussions to allows the sustainability of the primary objective of these systems; obtain points and sharp blades, primarily rectilinear. These technical and technological evolutions go hand in hand in units XIX to XV of the EUP with the development of backs, and in particular of truncated backed points which will take an increasingly important place in the balance of tools, until occupying a central place in the typological corpus in layers XVI and XVII. Figure 6 shows the gradual development of these tools which will represent more than 20 to 40% of the EUP tools before clearly collapsing starting at layer XIV, the last unit of the Northern Early Ahmarian of Ksar Akil. Figure 6 also balances the 1937-1938 London collections based on Ohnuma's data [101] and my own analyses of the 1947-1948 Harvard collections. The two graphs show a clear concordance between these two collections, with a bell shape and backs that only develop in the Early Ahmarian and in an extremely marginal way (less than 1 to 2% of the tools) in the units of the IUP before disappearing after units XV-XVI. This development curve remarkably matches the bipolar debitage development curve, which gradually increases in strength across all of the Early Ahmarian units before collapsing after layer XV. There is a pertinent correlation here because a large part of these secondary appointments by abrupt retouching and direct truncations has the effect of taking up the slight curvature of these pointed blades in order to obtain points with a rectilinear profile. Here we can discern an equilibrium of the technical system whose production objectives remain identical; obtain laminar blanks that are both sharp and flat. While the change hard hammerstone percussion to soft hammerstone percussion allows greater flexibility in debitage, as a side effect, it also mechanically induces, a modification of the general morphology of the desired blanks. This side effect of percussion techniques affects the systems for which the structure of debitage targets, but nevertheless remains immutable over time; obtaining slender, pointed, and rectilinear supports. The system is balanced here by recourse to the opening of opposing striking planes on the cores and by the implementation of an abrupt secondary retouching, shortening the blank to rectify the curvature while preserving, or amplifying its acuminate character. Within the stratigraphy, the parallelism within the progressive development of these two technical indicators; modifications of percussion techniques, so, the development of bipolarity, go hand in hand in this laminar universe with a progression of the place given, within the tools, to the points with backed backs.

**Supplementary Note 5: After the EUP of Ksar Akil**

The archaeological record of Ksar Akil is still very rich after layer XIV. My technical quantifications stopped at layer XIII, and were followed by a schematic analysis of all the overlying units. However, it was fundamental to integrate layer XIII in this quantified analysis because it technically shows the complete overturning of the systems of the Early Ahmarian of Ksar Akil in a very distinct technical sphere. Once again, this changeover is not abrupt and one can see in both the bipolarity indicators and in the typological indicators that layers XV and XIV already initiated an overturning of the technical systems, which only appears fully completed in layer XIII. Layer XIII is generally not attributed to a specific phase in the analyses of Ksar Akil's collection. This would then be phase 3 of Williams and Bergman, whose figure 1 shows that the phase is generally considered without lithic industry (collections 1937-1938?) or as an undetermined Upper Paleolithic ("unnamed UP", "unassigned"), perhaps also because of the rarity of typological tools in this unit; 42 tools including 23 burins on truncation and 12 scrapers. However, this unit is composed of 547 lithic pieces that are technically well diagnosed as to the precise functioning of this technical system. The debitage is now entirely undertaken using organic soft percussion, a trend that began as early as layer XVIII on the lightest laminar blanks. The industry from layer XIII is light, essentially lamellar, and shows the ability to achieve remarkably standardized lamellar blanks. The notion of predetermination, as understood in the preliminary phases of the IUP and EUP remains present, but these very slender micropoints are easily confused with acute bladelets presenting a morphology of great regularity. This technical atmosphere of highly standardized rectilinear bladelets resulting from converging unipolar debitages is found in the overlying levels XII, XI, and X, associated with fine regular direct, inverse, or alternate, retouches, of which several are attested from this layer XIII. In 2017, Christopher Bergman finally identified affinities between these overlying industries of his Phase 4 (Fig. 1) and the Southern Early

Ahmarian ensembles [79], a proposal however that concerned only layer XI and overlying layers[‡]. More than affinities, we can consider here that these industries are registered in Ksar Akil from phase 3, and not from phase 4. Strictly speaking, these industries correspond to the industries of the Early Ahmarian but such as they were recognized in the south-Levantine zone starting from the beginning of the 1980s. Jim Phillips would finally find his offspring from the region of Abu Noshra… I attribute layer XIII to the Protoaurignacian (Fig. 1).

**Supplementary Note 6: From East and West. Back to Mandrin, downgrading, reclassification, pieces of the puzzle of Western Europe**

Even though the divergences in the correlations between East and West had been recognized since the 1980s, the study by Kadowaki *et al*. [47] is historiographically interesting because it is the first to really reintroduce the divergences, technically very marked, between the two entities attached to the Early Ahmarian on one hand, and the Protoaurignacian on the other. It should be noted that Kadowaki *et al*. in their study compare the European Protoaurignacian to phase 4 of Ksar Akil as defined by Williams and Bergman [37], leading them to conclude that the Protoaurignacian may have anteriority over its Levantine counterparts, reintroducing Paul Mellars' proposal from ten years earlier [29]. From the point of view of the general technical structure of these industries, however, phase 4 of Ksar Akil (layers IX-XI) does not represent the oldest industries of Ksar Akil that are technically comparable to the Protoaurignacian, which I place as early as layer XIII. The analysis of the collections suggests that connections with the European Protoaurignacian should be sought in the units stratigraphically underlying Williams and Bergman's phase 3; these units record a collapse of bipolar debitage (layers XV-XIV), followed by a complete shift in debitage toward unipolarity. Layer XIII, wholly oriented toward obtaining

[‡]"This view is essentially in agreement with the suggestion by Mellars (2006a:172-175) for the similarity between the Protoaurignacian (or the Fumanian) and the assemblages from Boker A (the southern Early Ahmarian) and Ksar Akil Levels XI-IX (mainly Phase 4) that are commonly characterized by small, lightly retouched bladelet points of Font Yves and el-Wad types" (Mellars 2006b).

pointed and rectilinear bladelets with convergent unipolar debitage, is part of the Protoaurignacian phylum, in its strict sense, as recognized in continental Europe.

Here the Levant/Europe correlation is easiest to establish. In my opinion the correlation between the Early Ahmarian of Ksar Akil [and more broadly of the entire northern Levantine zone) and the Protoaurignacian must be definitively abandoned. The technical proximity of the European Protoaurignacian to the industries of Ksar Akil can be documented in more recent strata, emerging with Layer XV and becoming evident in the overlying units XIV and XIII.

One should be surprised by the systematic nature of the confusion established during the 17 years between NEA and SEA on this issue of the Levantine origins of the Protoaurignacian. In this international debate, the confusion has astonishingly led to no longer taking account the specific technical characteristic of the Ahmarian of Ksar Akil, although richly documented graphically in the works of Azoury and Ohnuma.

If the correlations between the European Protoaurignacian and phase 4 of Ksar Akil introduced by Kadowaki *et al*. [47] are in line with the initial proposals of Mellars [29], the correlations do not include the first units of Ksar Akil into this Protoaurignacian technical movement. The stratigraphic phasing shift that I highlight here indicates that in much older stratigraphic units- as early as layer XIII- at Ksar Akil it is possible to establish a strict technical community with the European Protoaurignacian. It is also possible to imply that the origin of these industries is clearly perceptible in even older units recognizable as early as in layer XV. On the basis of the technical realities at hand, this correlation allows us to contradict on different bases the question of a chronological anteriority of the European Protoaurignacian over the Levantine ensembles as proposed in Kadowaki *et al*. [47]. But much more than the correlations of Bayesian models, often rather fragile, it is also, and above all, the clear technical continuity between layer XIII and the underlying sets, and this in a much more general way until the very beginning of the IUP in layer XXV which induces, in a structural approach of this problem, the impossibility of an anteriority of the first European Protoaurignacian over any of the technically comparable forms documented at Ksar Akil[§].

**Supplementary Note 7: The Châtelperronian Question**

These technical categories of slender pieces whose backs are structurally quite disparate in origin represent the main technical decoders used to bring the Châtelperronian closer to certain Mousterian ones in the same geographic area [46, 66-68]. By recentering the question on elements that are technically and typologically comparable, i.e. backed points made on blades -issued from true blade debitages- to the exclusion of simple supports that are morphologically slender and of varied technological origin, one realizes that these precise technical categories are generally absent from the Mousterian industries and in no way characterize their technical framework. The question of Mousterian backed pieces by far transgresses not only the geographic distribution of the Châtelperronian [110] but also a chronological distribution restricted to only the recent phases of the Mousterian [8]. It is probably in older sets of the Mousterian (pre-MIS 3) that the distribution of true backed points, backed backs, and on blades would potentially be the most important [7]. At Mandrin, the concept of backed pieces - on any technical blank- can be documented throughout all the units of the sequence, from MIS 5 up until embodying the typologically dominant category of layer  D somewhere ~51 ka. It is not suggested here that there is any phyletic link between these industries of the Post-Neronian I of Mandrin D and the Châtelperronian, but simply points out that the links between Mousterian and Châtelperronian

[§]"several researchers (e.g., Bergman, 1988; Kuhn et al., 2003; Marks, 2003:255; Goring-Morris and Davidzon, 2006; Tsanova et al., 2012) have noted the regional variability between the southern arid zone (Negev, Sinai, and Jordan) and the northern Levant. Specifically, Goring-Morris and Davidzon (2006:106-107) suggest that the Ahmarian assemblages in the northern Mediterranean zone (including Üçagızlı B, B4-1 and C, Ksar Akil XX-XV, Yabrud II, Kebara IV-III, and Qafzeh E-D) are characterized by the use of a bi-directional blade/bladelet knapping method in addition to a single platform core reduction that is predominant in the southern Ahmarian assemblages. Goring-Morris and Davidzon (2006) also mention that blades/bladelets removed from opposed-platform cores in the northern Ahmarian appear to be relatively straight and robust in comparison with the slender and pointed blades/bladelets from single-platform narrow-fronted cores in the southern Ahmarian. Such differences in blank forms are related to the variability in the size and form of points and their retouch patterns in the Ahmarian assemblages. This aspect has been examined through the definition of various point types (e.g., el-Wad points, Ksar Akil points, and pointes à face plane; Bergman, 1981) that have been noted to occur in different frequencies between the southern and northern Ahmarian assemblages (Kuhn et al., 2003)" (Kadowaki et al. 2015).

cannot be structured on the basis of generic technical categories such as the presence of more or less slender blanks and presenting any category of backs, even if they were natural or modified by some marginal abrupt retouching. The recent modification of the Châtelperronian's distribution area, which now extends to the Iberian Mediterranean coast [76], calls into question the geographic correlation established between these generic Mousterian backs and the Châtelperronian. The question of Ruebens et al. [68] then poses a problem if we consider, as these authors, this vision of backed points which considers them "a technology absent in early modern human technologies outside of South Africa" [68][**]. In this contextualization, the backed points *stricto sensu*, made on blades and not on elongated blanks, and actually backed by truncation, are either anecdotal or simply absent from the Mousterian series in the very range of the Châtelperronian and occupy, more generally, a position that is at best marginal within the MP industries of continental Europe.

Such a precise and trans-Mediterranean technical connection model has direct implications on the question of the biological author of the Châtelperronian. If we consider that some of the phases of the Early Ahmarian of Ksar Akil and the European Châtelperronian are products of the same technical tradition, is it then possible to envisage that this tradition could be shared by two distinct humanities? Unless we consider very powerful processes of acculturation, this hypothesis immediately appears quite fragile. Since the discussions around Saint-Cesaire [63], only the sequence of la Grotte du Renne in Arcy-sur-Cure would make it possible to recognize the population at the origin of the Châtelperronian. The study by Welker *et al*. in 2016, which combined biomolecular analyses with direct dating on human remains at Arcy, seems to close the debate in favor of Neandertals [111]. Yet the assertion according to which there would be a clear distinction, within the genus *Homo*, of human clades based on ancient protein analysis, and

[**]"Especially, the recurrence of elongated and backed elements in both the late Mousterian and Châtelperronian in a restricted geographic region, and its near absence in other Mousterian, transitional, and IUP technocomplexes, seem to support an argument in favor of local continuity (…) The continuation of the technological concept of backing, absent elsewhere, seems to favor a local, Neanderthal origin for the Châtelperronian (…) If the Châtelperronian was (partially) made by modern humans, it is still to be determined from where these moderns came (…), and why they started making backed points, a technology absent in early modern human technologies outside of South Africa" (Ruebens et al. 2015).

which would make it possible to recognize the craftsman of the Châtelperronian from Arcy [111] should probably be abandoned [112]. This biomolecular analysis should therefore be considered inconclusive, in contrast to the classic anthropological analysis carried out at Arcy which, in turn, demonstrates the Neandertal characteristics of the human remains found in these units [113]. The questions raised around the integrity of these archaeological assemblages, which have shown through radiometric results, that a third of the bone elements from these Châtelperronian levels would be intrusive, remain unanswered [114, but see 115]. These questions may also refer to the 'Mousteroid' component of Arcy's Châtelperronian technical systems which would never be found in a collection in which no underlying Mousterian level has been recorded [59].

These questions must now be combined with the proposition that the Châtelperronian presents some very precise technical affinities with Ksar Akil's Northern Early Ahmarian, suggesting that Neanderthals may not have been at the origin of the Châtelperronian. This hypothesis is refutable and could therefore be confirmed or rejected either by the discovery of *H. sapiens* remains in the Châtelperronian assemblages or by the discovery of DNA testifying to a recent interbreeding between these populations. However, in this phase of contacts between populations on the European continent, we note that even though the presence of Neanderthal DNA in the first *H. sapiens* populations is well documented [116-120], the reverse is not true and that the genetic sequencing of the most recent Neanderthal populations in Europe shows the absence of any *H. sapiens* introgression within them [121]. If continuing paleogenetic analyses confirm such a pattern, even though the genetic flows between different populations now appear to be fairly systematic [122], the implications for the relationships between these two populations would be of central importance for understanding the precise historical and ethnographic interactions that existed between them at the time of Europe's colonization. Future data will have to question these propositions but, regarding technical systems, the Châtelperronian / NEA convergence makes it possible to posit that the origin of these technical traditions is to be sought in the Levantine space, therefore implying that the first carriers of this tradition in Europe may well be, in fact, the *H. sapiens* populations.
